# Stromal mitochondrial reprogramming and transfer reveal a chemotherapy-associated tumor state in ovarian cancer

**DOI:** 10.64898/2026.11.19.767697

**Authors:** Roja Baruwal, Paige Matusiak, Huda Atiya, Geyon Garcia, Angelina Li, Priscilla Tang, Swathi Suresh, Ben Johnson, Hui Shen, Alexandra Glathar, Anthony Cillo, Francesmary Modugno, Lauren Borho, Nadine Hempel, Leonard Frisbie, Lan Coffman

## Abstract

Mitochondrial transfer from stromal cells promotes ovarian cancer progression, yet the functional state of donor mitochondria remains poorly understood. Here we define a stress-adapted mitochondrial state in carcinoma-associated mesenchymal stem cells (CA-MSCs), characterized by increased respiratory and glycolytic activity despite increased mitochondrial proton leak and inefficient coupling of respiration to ATP production, oxidative stress, and elevated mitochondrial DNA variant burden. We identify WT1 as a regulator of mitochondrial reprogramming and PDK4 as a metabolic adaptation critical for CA-MSC growth. We further show that CA-MSCs preferentially transfer highly oxidized mitochondria to ovarian cancer cells. In a freshly-dissociated primary high grade serous ovarian cancer patient tumor, using mitochondrial variant tracking, we identify a transcriptionally distinct tumor population sharing mitochondrial variant patterns with stroma, in line with stromal–tumor mitochondrial transfer. This transcriptional signature is enriched in residual tumors following chemotherapy across four independent patient cohorts. Together, these findings establish stromal mitochondrial reprogramming and the preferential transfer of oxidatively-stressed mitochondria as important features of ovarian cancer heterogeneity and persistence following chemotherapy.

## Introduction

High-grade serous ovarian cancer (HGSOC) is one of the most common and lethal gynecologic malignancies. It is characterized by early dissemination, extensive intratumoral heterogeneity, and frequent resistance to standard therapies^1,2^. Although aerobic glycolysis has long been recognized as a prominent feature of ovarian cancer metabolism, increasing evidence demonstrates that mitochondrial metabolism also contributes to ovarian tumor growth and metastatic progression and influences response to chemotherapy^3,4–6,7,8^. Beyond ATP production, mitochondria function as central signaling organelles that regulate redox balance, metabolic plasticity, and cell fate decisions^9,10^. Accordingly, alterations in mitochondrial respiratory activity, redox balance, and fusion–fission dynamics are increasingly recognized as important features of ovarian cancer^3,11^. However, most studies of mitochondrial remodeling in ovarian cancer have focused on malignant cells, despite the critical role of the tumor microenvironment in ovarian cancer progression.

Ovarian cancer progression is critically dependent on continuous interactions with the tumor microenvironment, which includes immune cells, vasculature, extracellular matrix, and diverse stromal populations^1^. Ovarian cancer can also remodel mitochondrial function within surrounding non-malignant cells. For example, ovarian cancer can facilitate immune evasion by disrupting mitochondrial metabolism in T cells^12,13,14^. These findings highlight mitochondrial remodeling of the tumor microenvironment as a potential pro-tumorigenic process. However, mitochondrial remodeling within the stromal compartment of ovarian tumors remains comparatively understudied. One of the stromal cell populations that we and others have found to play key roles in ovarian cancer are mesenchymal stromal/stem cells (MSC). MSCs are multipotent stromal progenitor cells found in most adult tissues^15^. However, we previously demonstrated that ovarian cancer cells reprogram MSCs into cancer-associated MSCs (CA-MSC), a functionally distinct stromal population that promotes tumor growth, immune suppression, treatment resistance, and metastatic progression^16–21^. This tumor-supportive phenotype distinguishes CA-MSC from nontumorigenic or normal MSC (nMSC)^16,21^. We further demonstrated that this reprogramming involves epigenomic alterations, including changes in DNA methylation, chromatin accessibility, and histone modifications, resulting in a partial mesenchymal-to-epithelial transition (MET) in CA-MSC, with Wilms Tumor 1 (WT1) playing a crucial role in this reprogramming process^21^. Whether this tumor-induced reprogramming extends to the mitochondrial and metabolic state of CA- MSC remains unclear.

The mitochondrial state of CA-MSCs is of particular interest, as CA-MSCs can directly transfer mitochondria to ovarian cancer cells^22^. We recently showed that CA-MSC donate mitochondria to ovarian cancer cells, increasing tumor cell respiration, proliferation, and chemoresistance while promoting intratumoral heterogeneity and metastatic progression^22^. Stromal-to-cancer mitochondrial transfer has also been observed across multiple malignancies, including breast, colorectal, brain, and hematologic cancers^23–26^. While the pro-tumorigenic consequences of stromal-to-cancer mitochondrial transfer are increasingly recognized^22–26^, the mitochondrial state of the donor CA-MSC population remains undefined. Specifically, it remains unknown how CA- MSC mitochondria differ from nMSC, how these differences arise during tumor-induced reprogramming, and whether they influence the mitochondria transferred to ovarian cancer cells. Defining the mitochondrial and metabolic state of CA-MSC may therefore provide further insight into how tumor-induced stromal reprogramming influences mitochondrial transfer and ovarian cancer progression.

Here we define a distinct mitochondrial and metabolic state in CA-MSC characterized by increased bioenergetic activity, despite mitochondrial inefficiency and oxidative stress. Furthermore, we find this mitochondrial state is accompanied by structural, genomic, and transcriptional remodeling. We identify WT1 as a regulator of this mitochondrial phenotype, with PDK4 contributing to the glycolytic component of CA- MSC metabolic reprogramming. Importantly, we demonstrate that oxidatively stressed CA-MSC mitochondria are preferentially transferred to ovarian cancer cells. We further identify a tumor subpopulation within primary patient-derived high grade serous ovarian cancer (HGSOC) that shares mitochondria with CA-MSCs, confirming the relevance of mitochondrial transfer in human disease. Moreover, this subpopulation has a distinct transcriptional profile indicating unique features compared to non-mitochondrial recipient tumor cells and this subpopulation signature was significantly enriched in residual tumors collected after neoadjuvant chemotherapy (NACT) compared with matched pre-NACT tumors across four independent patient cohorts. Together, these findings establish the development of mitochondrial dysfunction as a critical component of the CA-MSC phenotype with the selective donation of these dysfunctional mitochondria to tumor cells which promote unique tumor characteristics including therapy resistance.

## Result

### CA-MSC exhibit a distinct bioenergetic profile

We previously demonstrated that CA-MSC transfer mitochondria to ovarian cancer cells^22^, yet whether CA-MSC themselves exhibit distinct mitochondrial functional properties compared with nMSC remains unclear. To address this, we performed extracellular flux analysis using the Cell Mito Stress Test in primary patient-derived nMSC and CA-MSC (Fig1. **A–G**). CA-MSC were isolated from cancer involving the fallopian tube of patients with HGSOC, whereas nMSC were isolated from fallopian tube tissue of patients undergoing surgery for benign indications. Both MSC populations were isolated and validated using our established protocols^17,21^. CA-MSC exhibited increased mitochondrial respiration (Fig. **1A**), with a trend towards higher basal oxygen consumption rates (OCR; Fig. **1B**) and significantly higher maximal respiration (Fig. **1C**).

**Figure 1.**
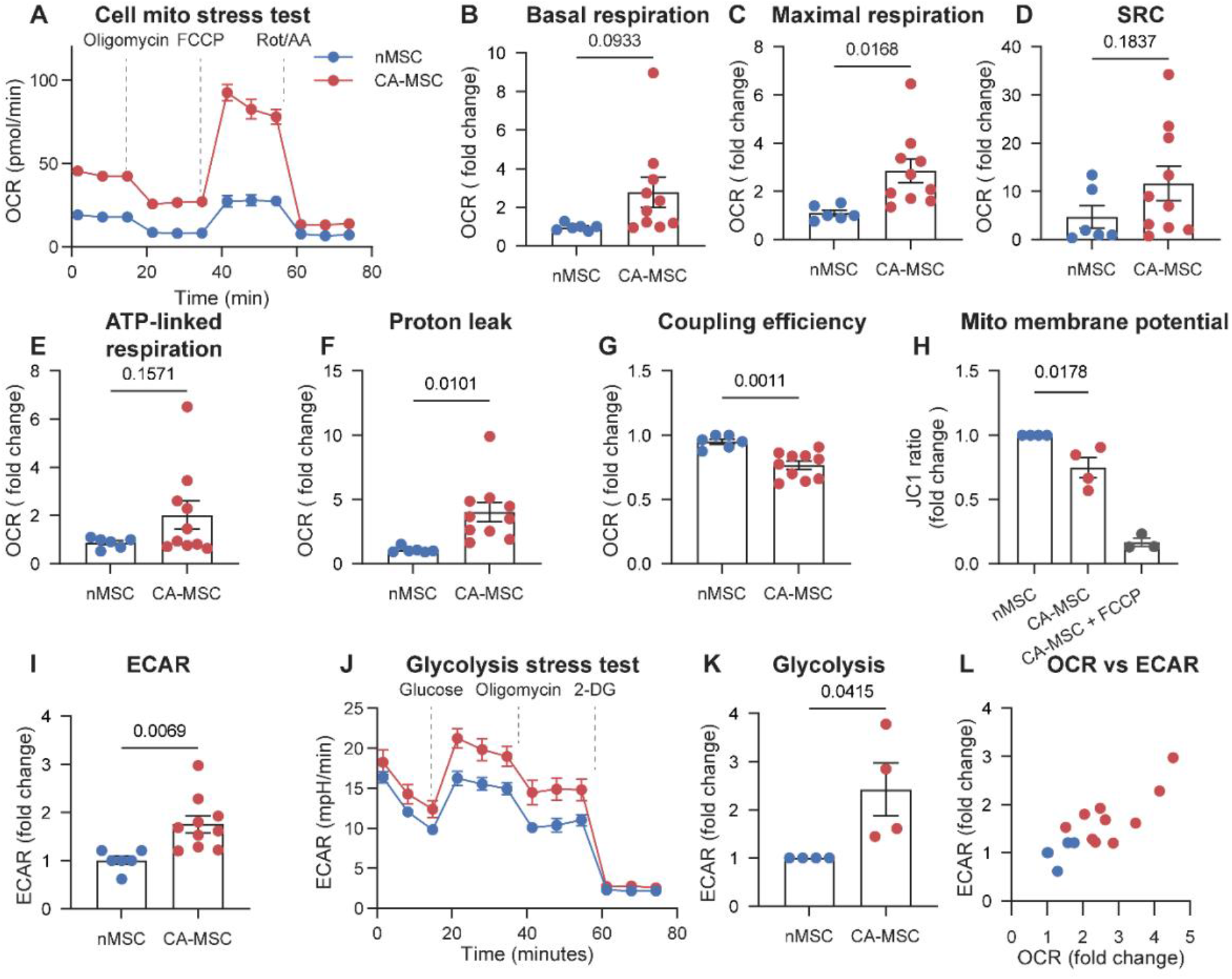
| CA-MSC exhibit a distinct bioenergetic profile. **A,** Representative oxygen consumption rate (OCR) profiles from Cell Mito Stress Tests of patient-derived nMSC and CA-MSC following sequential injection of oligomycin, carbonyl cyanide 4-(trifluoromethoxy)phenylhydrazone (FCCP), and rotenone/antimycin A. **B–G,** Quantification of basal respiration (B), maximal respiration (C), spare respiratory capacity (D), ATP-linked respiration (E), proton leak (F), and coupling efficiency (G) (n>3). **H,** Mitochondrial membrane potential assessed by JC-1 fluorescence ratio, with FCCP-treated CA-MSC included as a depolarization control (n=4). **I,** Extracellular acidification rate (ECAR) quantification from cell mito stress test. **J,** Extracellular acidification rate (ECAR) profiles from Glycolysis Stress Tests following sequential injection of glucose, oligomycin, and 2-deoxyglucose (2-DG). **K,** Quantification of glycolysis (n=4). **L,** Relationship between OCR and ECAR in nMSC and CA-MSC, expressed as fold change relative to nMSC. Quantitative measurements are expressed as fold change relative to nMSC, except for the OCR and ECAR profiles in (A) and (J). Data are presented as mean ± sem. Statistical significance was determined using unpaired t-test, with exact p values indicated in the graphs.

Spare respiratory capacity (SRC) was also higher on average in CA-MSC, although this difference did not reach statistical significance (Fig. **1D**). Despite increased oxygen consumption, ATP-linked OCR was not proportionately elevated (Fig. **1E**), which is further illustrated by a significantly reduced coupling efficiency (Fig. **1G**) and increased proton leak (Fig. **1F**). Consistent with increased proton leak, CA-MSC also exhibited significantly reduced mitochondrial membrane potential, indicating partial mitochondrial depolarization relative to nMSC (Fig. **1H**). Simultaneous analysis of extracellular acidification rate (ECAR) suggested increased glycolysis in CA-MSC (Fig. **1I**). Further analysis using the Glycolysis Stress Test confirmed elevated glycolysis in CA-MSC relative to nMSC (Fig. **1J,K**). Together, these data indicate that CA-MSC adopt a more bioenergetic phenotype (Fig. **1L**), with increased glycolysis and respiration, but with oxygen consumption that is partially uncoupled from ATP production, indicating a level of sub-optimal mitochondrial function.

## CA-MSC exhibit altered mitochondrial morphology

Given their inefficient mitochondrial respiration, we next examined whether bioenergetic alterations in CA-MSC were accompanied by changes in mitochondrial morphology. To assess this, nMSC and CA-MSC were labeled with COX8-GFP and imaged using confocal microscopy (Fig. **2A**). CA-MSC exhibited significant differences in mitochondrial morphology compared to nMSCs (Fig. **2A**). Specifically, CA-MSCs had a higher mean mitochondrial aspect ratio (Fig. **2B**) and form factor (Fig. **2C**) compared with nMSC, along with increased branch length per mitochondrion (Fig. **2D**), indicating a shift toward more elongated mitochondrial structures. The number of branches per mitochondrion (Fig. **2E**) was also increased, suggesting greater mitochondrial network branching.

**Figure 2.**
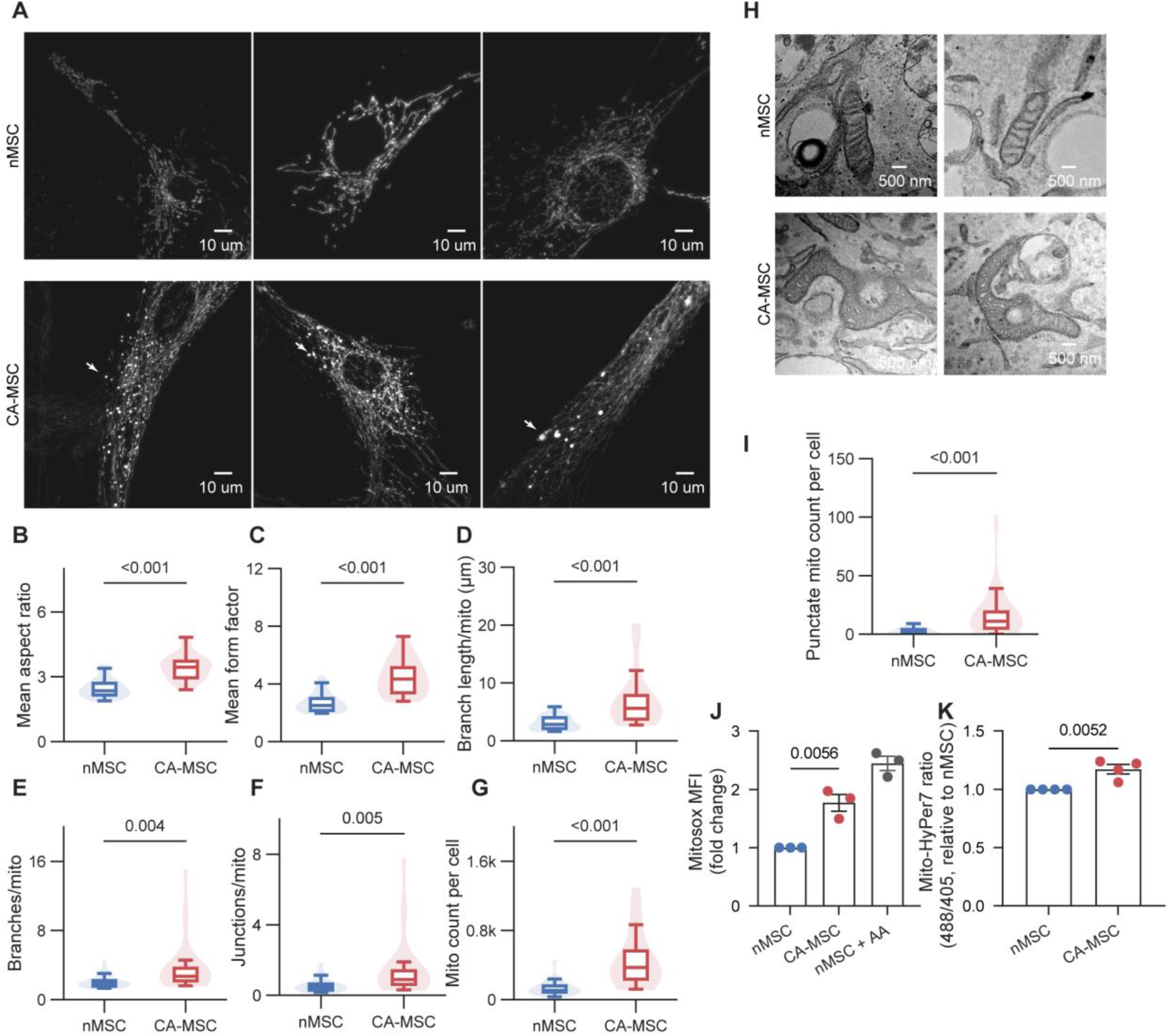
| CA-MSC exhibit altered mitochondrial morphology and elevated mitochondrial oxidative stress. **A,** Representative confocal images of mitochondria in patient-derived normal MSC (nMSC) and carcinoma-associated MSC (CA-MSC) expressing mitochondrially targeted COX8-GFP. Images were converted to grayscale for visualization. White arrows indicate punctate mitochondrial structures in CA-MSC. Scale bars, 10 μm. **B–G,** Quantification of mitochondrial morphology using the Mitochondria-Analyzer ImageJ plugin, including mean aspect ratio (B), mean form factor (C), branch length per mitochondrion (D), branches per mitochondrion (E), total mitochondrial area per cell (F), and total mitochondrial number per cell (G). **H,** Representative transmission electron microscopy (TEM) images of nMSC and CA-MSC showing donut-shaped mitochondria in CA-MSC. Scale bars, 500 nm. **I,** Quantification of punctate mitochondrial structures per cell from confocal images. **J,** Flow cytometric quantification of mitochondrial superoxide using MitoSOX Red. Antimycin A (AA) was used as a positive control. **K,** Flow cytometric quantification of mitochondrial hydrogen peroxide-associated oxidation using mitochondrially targeted HyPer7. Statistical significance was determined using two- tailed unpaired Student’s t-tests. For mitomorphology analysis (B-G and I), data are shown from one representative nMSC and CA-MSC line comparison. Two additional line-pair comparisons showed differences in the same direction and are presented in Supplementary Fig. 1A,B. For J,K, each dot represents a different patient line. Exact *P* values are indicated in the graphs.

Visual inspection revealed punctate mitochondria interspersed within the tubular mitochondrial network in CA-MSC (Fig. **2A**). Quantification demonstrated an increased number of these punctate mitochondria per cell in CA-MSC compared with nMSC (Fig. **2I**). CA-MSC also exhibited an overall increase in total mitochondrial area and mitochondrial count per cell (Fig. **2F,G**). Similar increases in mitochondrial aspect ratio, form factor, branching, mitochondrial number, and punctate mitochondrial structures were observed in two additional independent nMSC and CA-MSC line comparisons (Fig. **S1A,B**). At higher magnification, ring- and donut-shaped mitochondrial structures were observed by fluorescence microscopy (Fig. **S1C**) and transmission electron microscopy (Fig. **2H**). These structures were also observed in CA-MSC when staining with MitoTracker Deep Red (Fig. **S1D**). Together, these findings demonstrate heterogeneous mitochondrial remodeling in CA-MSC, characterized by the coexistence of elongated mitochondria and increased punctate mitochondrial structures.

## CA-MSC exhibit elevated mitochondrial oxidative stress

Since we demonstrate CA-MSCs have increased mitochondrial proton leak and altered mitochondrial morphology, both features associated with mitochondrial oxidative stress^27–30^, we next examined whether CA-MSC mitochondria also have elevated oxidative stress. CA-MSC showed significantly higher oxidation of the superoxide- sensitive probe MitoSOX (Fig. **2J**). Similarly, an increase in the oxidized to reduced form of the genetically- expressed hydrogen peroxide sensor HyPer7^31^ targeted to the mitochondrial matrix was observed in CA-MSC compared to nMSC (Fig. **2K**). Together, these findings demonstrate that many CA-MSC mitochondria exist in a stressed state, characterized by elevated oxidative burden.

## CA-MSC exhibit increased mitochondrial DNA mutational burden and enrichment of homoplasmic variants

We next sought to understand the mechanism of mitochondrial dysfunction in CA- MSCs. We first investigated if CA-MSCs were more enriched for mitochondrial genomic mutations. To quantify the mitochondrial DNA (mtDNA) variant landscape of patient- derived CA-MSCs, we first leveraged mitochondrial reads from previously-generated, bulk assay for transposase-accessible chromatin with sequencing (ATAC-seq) data^21^. In these bulk data, we found CA-MSC libraries had significantly more somatic mtDNA single-nucleotide variants (SNVs) than nMSC libraries (median 16 vs. 8 variants per library; Wilcoxon rank-sum test, W = 0, p = 0.029) (Fig. **3A,B**). Next, to further assess heterogeneity and variant prevalence at a per-cell level, we also performed single-cell ATAC-seq in CA-MSCs and nMSCs. Single-cell analysis revealed a trend towards more mtDNA SNVs per-cell in CA-MSC than in nMSC (2.66-fold increase; linear mixed-effects model on log10-transformed per-cell counts, donor as random effect; β = 0.43, SE = 0.16, t(3) = 2.64, p = 0.078) (Fig. **3D**). Across both single-cell and bulk ATAC-seq datasets, variants were detected in mtDNA features critical for mitochondrial translation and metabolic function; this included mitochondrial ribosomal genes (RNR1, RNR2), Complex I genes (ND4, ND5), Complex III gene CYTB, and the regulatory D-loop region (Fig. **3C**, Fig. **S2**). These findings support that CA-MSC harbor an increased mtDNA variant burden, including variants within functionally relevant mitochondrial genomic regions.

**Figure 3:**
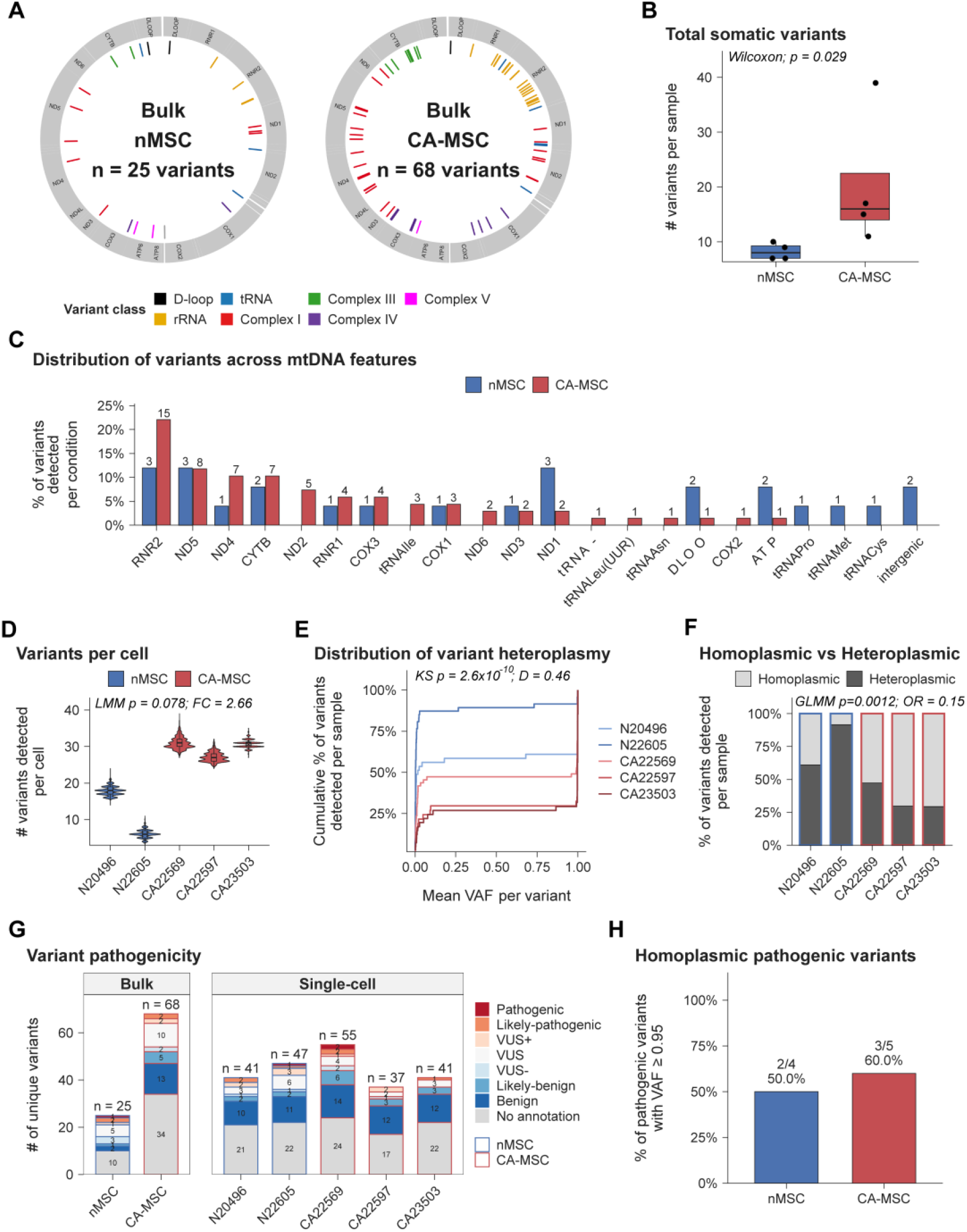
CA-MSC exhibit increased mitochondrial DNA mutations with enrichment in homoplasmic variants Mitochondrial variants detected in patient-derived CA-MSC and nMSC, across bulk and single-cell ATAC-seq samples. Bulk samples: n=4 normal vs n=4 HGSOC. Single-cell samples: n=2 normal vs n=3 HGSOC. **A,** Loci of somatic mitochondrial single- nucleotide variants (mtDNA SNVs) detected across bulk samples. Tick mark colors indicate mitochondrial feature designation based on variant locus. **B,** Quantified cross- condition comparison of total somatic SNV counts between bulk CA-MSC and nMSC libraries. **C,** Distribution of bulk sample somatic SNVs across mitochondrial features. Bar lengths indicate proportion of all mtDNA variants detected in a given feature. Proportions across all features sum to 100%, per condition. Variant counts in each feature are shown above each bar. **D,** mtDNA SNV counts, per-cell, across single-cell samples. **E,** Global distribution of mean variant allele frequencies (VAF) across single- cell samples. **F,** Proportion of heteroplasmic and homoplasmic variants detected in single-cell samples. Heteroplasmic variants had variance-to-mean variant allele frequency ratio (VMR) surpassing 0.01. **G,** MitImpact4 pathogenicity annotations of SNVs across bulk and single-cell samples. Total variant count detected per sample are listed above each bar. (VUS: variant of unknown significance, VUS-: low-scoring VUS. VUS+: High-scoring VUS). **H,** Proportion of likely-pathogenic or pathogenic variants in single-cell samples achieving homoplasmy, per condition. Variants with an observed maximum variant allele frequency of 0.95 (VAF=0.95) were considered to have achieved homoplasmy.

Because a single MSC contains many mitochondrial genome copies, we next assessed variant allele frequency (VAF) of individual mitochondrial SNVs across single cells. VAF is defined as the proportion of variant alleles found at a given mitochondrial genomic locus across all mitochondria of a cell. A homoplasmic variant is one that is shared amongst all mitochondria (VAF > 0.95), whereas a heteroplasmic variant is inconsistently present across mitochondrial genomes of a given cell. In our single-cell ATAC-seq data, we observed a significant difference in the global distribution of variant allele frequencies between CA-MSC and nMSC (two-sample Kolmogorov–Smirnov test, D = 0.463, p = 2.64 × 10^-10^; n = 88 nMSC vs. 133 CA-MSC variants), with CA-MSC variants shifted toward higher VAF (Fig **3E**). Consistent with this, CA-MSC variants were less likely to be heteroplasmic than variants in nMSC (odds ratio = 0.15, 95% CI: 0.046–0.47; logistic mixed model with donor as random effect, z = −3.24, p = 0.0012) (Fig. **3F**); conversely, this indicates that a CA-MSC variant was more likely to be homoplasmic (i.e., shared amongst all mitochondrial genomes of a given cell) when compared to nMSC. These results suggest CA-MSC variants, when present, are more likely to be shared and fixed across mitochondrial genomes of a given cell.

To evaluate the potential functional impact of the mtDNA variants detected amongst nMSC and CA-MSC, we asked if any of our detected variants had known pathogenicity. Using MitImpact 4 pathogenicity annotation^32,^ we found most variants were unannotated or classified as benign (Fig. **3G****, Supplementary Table 2**). However, both nMSC and CA-MSC contained “pathogenic” and “likely pathogenic” mtDNA variants. In CA-MSC, pathogenic/likely pathogenic variants were found in ND1 (3715G>A), tRNA^Met^ (4428G>A; 4429G>A), ND5 (13288G>A; 13340T>C; 13916G>A), and CYTB (15414A>G). In nMSC, pathogenic/likely pathogenic variants were detected in ND1 (3460G>A, 3916G>A, 3779G>A, and 4103T>G), tRNA^Met^ (4435A>G), ND4 (11535A>G), and tRNA^Pro^ (16023G>A).

A variant’s heteroplasmy – the degree to which it is shared across mtDNA genomes – is an important determinant for the emergence of downstream mitochondrial dysfunction. Therefore, we next checked if any of the pathogenic or likely pathogenic-annotated variants achieved homoplasmy in our single-cell samples. In CA-MSC, three of five (60%) of these pathogenic/likely pathogenic variants achieved homoplasmy (VAF>0.95), compared to 2/4 (50%) in nMSC (Fig. 3H); in CA-MSC, the homoplasmic variants were found in genes essential for respiratory chain Complex I function – namely ND1 (3715G>A) and ND5 (13288G>A and 13916G>A) – whereas in nMSC, these homoplasmy-achieving variants were found in ND4 (11535A>G) and tRNA^Pro^(16023G>A).

Collectively, these results demonstrate that CA-MSCs harbor an increase in mitochondrial mutational burden, with enrichment of homoplasmic variants in functionally important mitochondrial loci.

## CA-MSC enriched for mitochondrial DNA variants exhibit complex-specific patterns of respiratory dysfunction

Given the enrichment of potentially pathogenic mitochondrial variants in CA-MSC, including variants within genes encoding electron transport chain (ETC) components, we next asked whether these mitochondrial genomic alterations were accompanied by changes in ETC composition and respiratory function. Because the functional consequences of individual mitochondrial variants cannot be readily assessed through genetic manipulation, we examined the same patient-derived CA-MSC lines used for single cell ATAC-seq, with the corresponding nMSC lines serving as controls, thus the mitochondrial genomic variants in each cell line tested below are known.

We first assessed the abundance of representative proteins from individual oxidative phosphorylation (OXPHOS) complexes relative to the mitochondrial marker TOMM20. NDUFB8 (Complex I), SDHB (Complex II), and COXII (Complex IV) showed increased abundance in two of the three CA-MSC lines examined, whereas the third line showed no appreciable differences relative to nMSC. UQCRC2 (Complex III) and ATP5A (Complex V) remained relatively unchanged (Supplementary Fig.**S3A,B**). These findings suggest interpatient variability in ETC protein abundance among CA-MSC.

We next performed Seahorse analysis on isolated mitochondria from nMSC and CA- MSC lines with characterized mitochondrial variant profiles, using substrates that support respiration through different ETC entry points. Mitochondrial variant analysis identified a likely pathogenic variant in MT-ND1 (3715G>A) in CA-MSC, which encodes a core subunit of Complex I involved in proton translocation. In contrast, no pathogenic or likely pathogenic variants in any ETC-encoding genes were identified in nMSC.

Under Complex I-supported conditions using pyruvate and malate, overall respiratory activity was largely maintained in CA-MSC mitochondria (Fig. **4A**). Notably, CA-MSC exhibited increased proton leak under these conditions, raising the possibility that the MT-ND1 (3715G>A) variant contributes to altered respiratory coupling during Complex I- supported respiration. Respiratory control ratio (RCR), an indicator of mitochondrial respiratory coupling, was also lower in CA-MSC compared with nMSC (Fig. **4A**). To expand the biological replicates, we performed additional Complex I-supported Seahorse analyses using independent nMSC and CA-MSC lines for which mitochondrial variant profiles were not available. Similar increases in proton leak were observed in CA-MSC, while RCR was not significantly altered **(Fig. 4D).**

**Figure 4.**
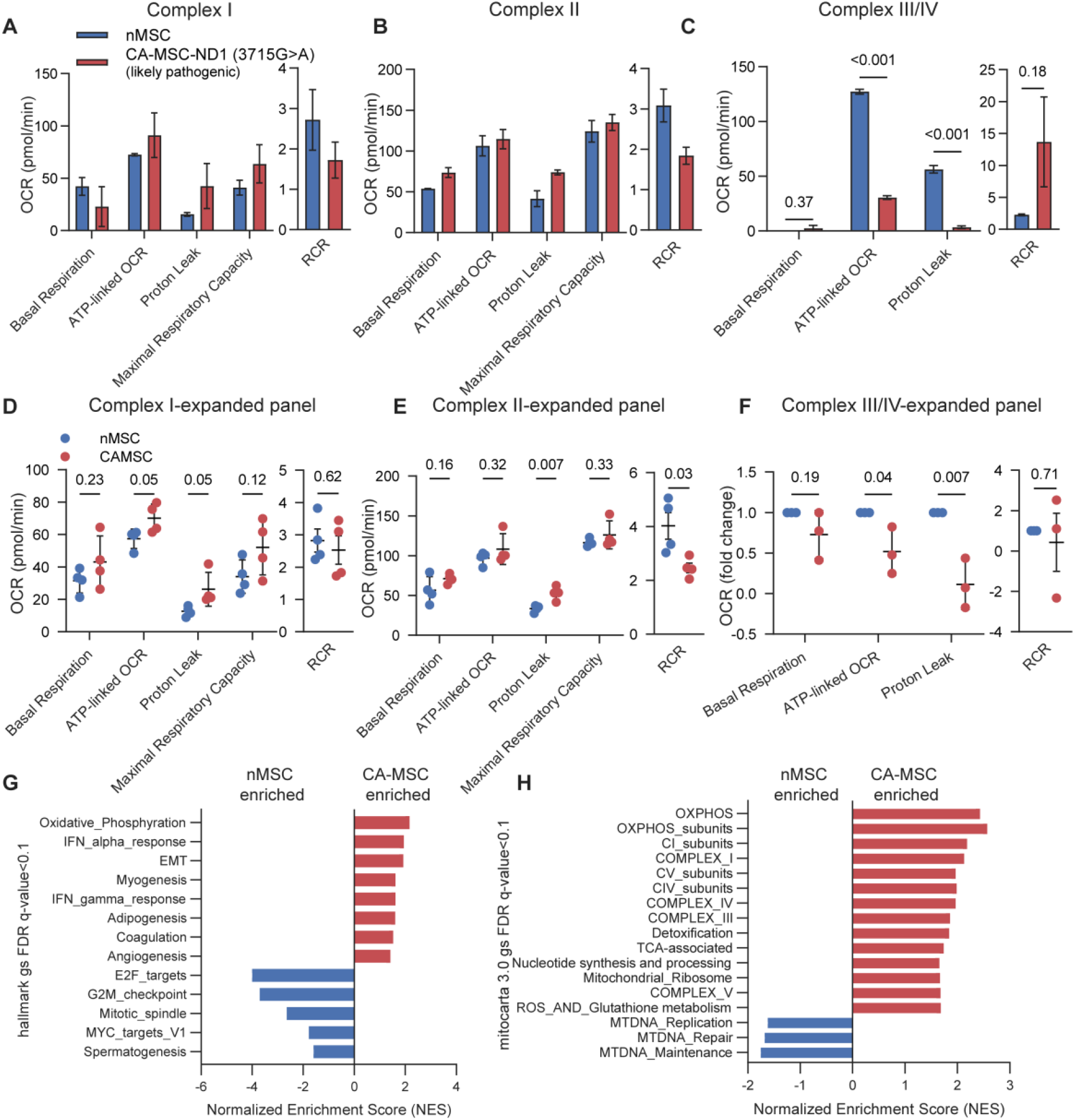
| CA-MSC exhibit complex-specific alterations in mitochondrial respiration and enrichment of oxidative metabolic pathways. **A–C,** Quantification of isolated mitochondrial respiratory parameters under Complex I- supported (A), Complex II-supported (B), and Complex III/IV-supported (C) conditions, including basal respiration, ATP-linked oxygen consumption rate (OCR), proton leak, maximal respiratory capacity (where measured), and respiratory control ratio (RCR). Complex I-supported respiration was assessed using pyruvate and malate as substrates. Complex II-supported respiration was assessed using succinate in the presence of rotenone to inhibit Complex I and prevent reverse electron transport. Complex III/IV-supported respiration was assessed using duroquinol as an electron donor to Complex III, bypassing Complexes I and II. **D–F,** Expanded analysis of Complex I-supported (D), Complex II-supported (E), and Complex III/IV-supported (F) mitochondrial respiration in additional independent patient-derived normal MSC (nMSC) and carcinoma-associated MSC (CA-MSC) lines. **G,** Hallmark gene set enrichment analysis (GSEA) of bulk RNA sequencing comparing nMSC and CA-MSC, showing pathways enriched in CA-MSC and nMSC. **H,** MitoCarta 3.0 pathway enrichment analysis of bulk RNA sequencing comparing nMSC and CA-MSC, showing enrichment of oxidative phosphorylation and respiratory complex-associated pathways in CA-MSC and mitochondrial DNA maintenance, repair, and replication pathways in nMSC. For A–C, isolated mitochondrial respiration was assessed using one nMSC and one CA- MSC line previously analyzed by mitochondrial scATAC-seq, with two replicates per condition (A,B) and three replicates per condition (C). The CA-MSC line harbored a likely pathogenic MT-ND1 (3715G>A) variant. For D–F, additional independent patient- derived MSC lines were analyzed to expand the respiratory assessment. Mitochondrial variant profiles were not available for these additional lines. Each data point represents one independent patient-derived MSC line. Data are presented as mean ± s.e.m. Statistical significance for D–F was determined using an unpaired *t*-test, with exact *P* values indicated in the graphs. For G,H, normalized enrichment scores (NES) are shown for pathways meeting a false discovery rate (FDR) threshold of *q* < 0.1.

Under Complex II-supported conditions using succinate in the presence of rotenone to inhibit Complex I and limit reverse electron transport, overall respiratory activity was largely maintained, while proton leak was higher and RCR was lower in CA-MSC mitochondria (Fig. **4B**). These findings were further supported by an expanded cohort of independent nMSC and CA-MSC lines, which demonstrated significantly increased proton leak and reduced RCR in CA-MSC **(Fig. 4E).**

We next assessed downstream Complex III/IV-supported respiration using duroquinol, which donates electrons directly to Complex III, bypassing Complexes I and II in the same MSC pair. CA-MSC mitochondria exhibited reduced ATP-linked respiration and proton leak compared with nMSC, whereas RCR was increased in CA-MSC (Fig. **4C**). Similar reductions in ATP-linked respiration and proton leak were observed in an expanded cohort of independent nMSC, n=3 and CA-MSC, n=3 lines (Fig. **4F**).

Together, these findings reveal distinct complex-specific patterns of ETC function in CA- MSC, characterized by largely maintained Complex I/II-supported respiration with increased proton leak and reduced downstream Complex III/IV-supported respiratory activity. The reduced respiratory activity following direct electron donation to Complex III suggests a potential functional limitation at Complex III and/or IV. Furthermore, the presence of a likely pathogenic MT-ND1 (3715G>A) variant in the CA-MSC line provides a potential genetic context for the altered Complex I-supported respiratory coupling, although its direct contribution remains to be established.

## CA-MSC exhibit transcriptomic reprogramming consistent with a stress-adapted mitochondrial phenotype

To determine whether the mitochondrial alterations observed in CA-MSC were accompanied by broader transcriptional reprogramming, we analyzed bulk RNA sequencing (RNA-seq) data from 6 patient-derived nMSC and 13 CA-MSC. Hallmark pathway analysis identified oxidative phosphorylation as the most strongly enriched program in CA-MSC, alongside broader enrichment of interferon response, epithelial– mesenchymal transition, adipogenesis, and angiogenesis programs (Fig. **4G**). In contrast, Hallmark pathways enriched in nMSC were not predominantly associated with mitochondrial or metabolic processes (Fig. **4G**).

We next focused specifically on mitochondrial pathways using MitoCarta gene sets^33^. CA-MSC exhibited broad enrichment of pathways related to OXPHOS complexes, tricarboxylic acid (TCA) cycle metabolism and ROS/glutathione metabolism (Fig. **4H**). In contrast, nMSC were enriched for pathways involved in mitochondrial DNA maintenance, repair and replication (Fig. **4H**). Together, these findings demonstrate that the functional mitochondrial phenotype of CA-MSC is accompanied by coordinated transcriptional reprogramming toward increased oxidative metabolism and mitochondrial oxidative stress.

## Tumor microenvironmental stress progressively drives the CA-MSC oxidative mitochondrial phenotype

We previously demonstrated that five days of coculture with ovarian cancer cells can induce CA-MSC-like changes in nMSC, a process termed cancer stimulation^17,21^. We therefore asked whether cancer stimulation also induces the oxidative mitochondrial phenotype observed in patient-derived CA-MSC.

To assess whether tumor-derived signals and microenvironmental stress contribute to mitochondrial oxidation, we exposed mitoHyPer7-expressing nMSC to adherent culture, adherent coculture with ovarian cancer cells, serum-free ultra-low attachment (ULA) culture, or serum-free ULA coculture with ovarian cancer cells.

Mitochondrial oxidation progressively increased across these conditions. Adherent nMSC exhibited the lowest mitoHyPer7 ratio, whereas cancer stimulation under adherent conditions produced a modest but significant increase in mitochondrial oxidation. Serum-free ULA conditions further increased mitochondrial oxidation in nMSC, which was augmented by coculture with ovarian cancer cells, reaching levels comparable to CA-MSC (Fig. **5A**). Representative images of cancer-stimulated MSC (CS-MSC) also revealed bright, punctate mitoHyPer7-488-positive mitochondrial structures (Fig. **5B**). These observations indicate that both tumor-derived signals and microenvironmental stress contribute to the oxidative mitochondrial phenotype of CA- MSC.

**Figure 5.**
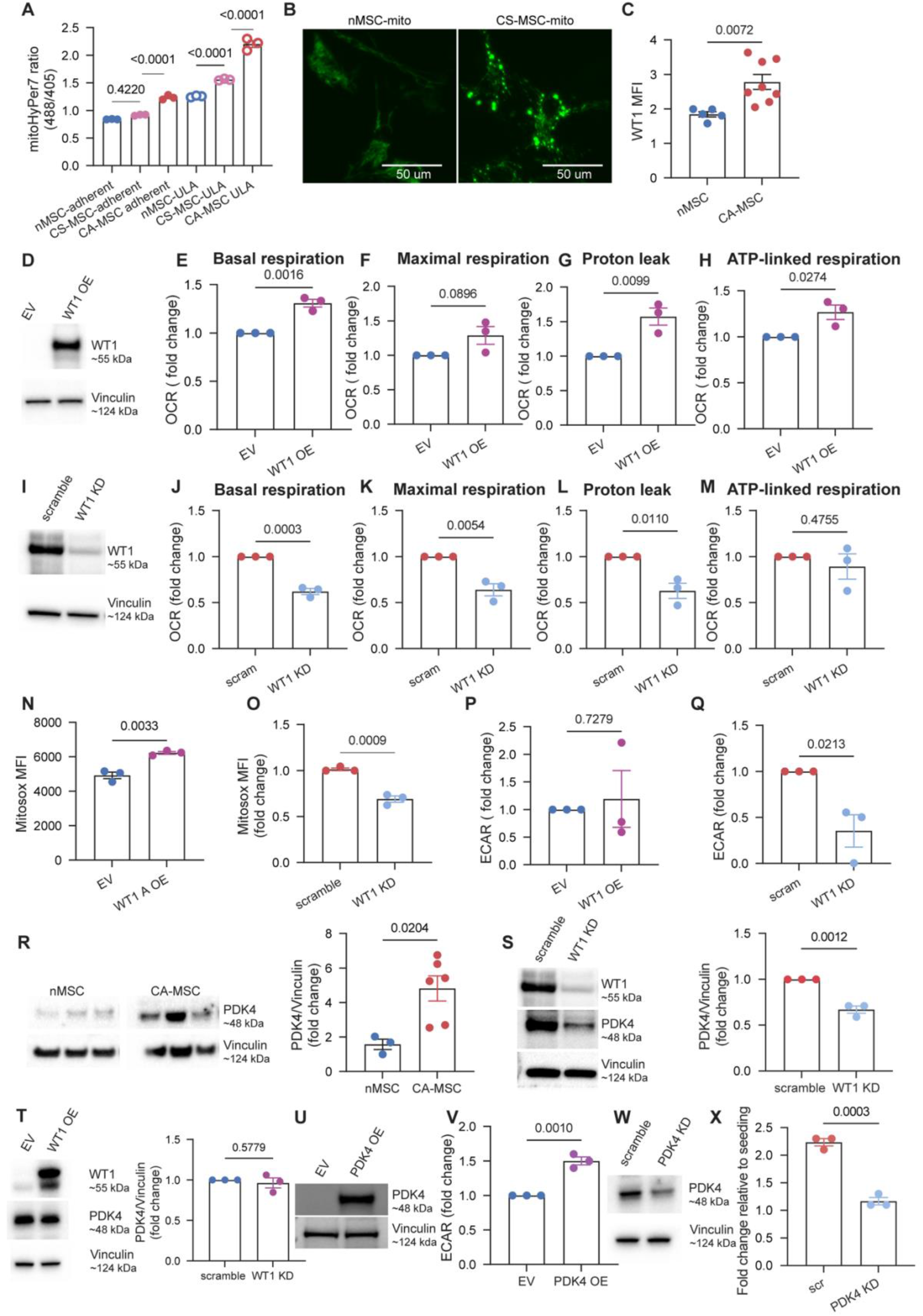
| Tumor microenvironmental stress and WT1 induction drives mitochondrial oxidative stress in CA-MSC whereas PDK4 contributes to the increased glycolytic respiration. **A,** Flow cytometric quantification of the mitochondrial HyPer7 oxidation ratio (488/405) in patient-derived normal MSCs (nMSC), carcinoma-associated MSCs (CA-MSC), and cancer-stimulated MSCs (CS-MSC). CS-MSC were generated by co-culturing nMSC with OVCAR3 cells for 5 days under adherent or ultra-low attachment (ULA) conditions (n = 3). **B,** Representative fluorescence images of mitochondrially targeted HyPer7 (488 nm channel) in nMSC and CS-MSC following 5 days of adherent cancer stimulation. Scale bars, 50 μm. **C,** Flow cytometric quantification of WT1 protein expression in nMSC and CA-MSC. **D,** Immunoblot validation of doxycycline-inducible WT1 overexpression in nMSC. **E–H,** Quantification of basal respiration (E), maximal respiration (F), proton leak (G), and ATP-linked respiration (H) following WT1 overexpression in nMSC. **I,** Immunoblot validation of WT1 knockdown in CA-MSC. **J–M,** Quantification of basal respiration (J), maximal respiration (K), proton leak (L), and ATP- linked respiration (M) following WT1 knockdown in CA-MSC. **N,O,** Flow cytometric quantification of mitochondrial superoxide using MitoSOX Red following WT1 overexpression in nMSC (N) and WT1 knockdown in CA-MSC (O). **P,Q,** Quantification of extracellular acidification rate (ECAR) following WT1 overexpression in nMSC (P) and WT1 knockdown in CA-MSC (Q). **R,** Immunoblot analysis and densitometric quantification of PDK4 protein expression in nMSC and CA-MSC. **S,T,** Immunoblot analysis and densitometric quantification of WT1 and PDK4 expression following WT1 knockdown in CA-MSC (S) and WT1 overexpression in nMSC (T). **U,** Immunoblot validation of PDK4 overexpression in nMSC. **V,** Quantification of ECAR following PDK4 overexpression in nMSC. **W,** Immunoblot validation of PDK4 knockdown in CA-MSC. **X,** Quantification of cell proliferation relative to initial seeding following PDK4 knockdown in CA-MSC compared with scramble controls.

## WT1 regulates mitochondrial and metabolic reprogramming in CA-MSC

We next sought to identify transcriptional regulators that may underlie this coordinated metabolic reprogramming. We previously demonstrated that the transcription factor Wilms tumor 1 (WT1) plays a key role in the epigenetic reprogramming of CA-MSC by ovarian cancer cells^21^. WT1, as expected, was elevated in patient-derived CA-MSC lines in comparison with nMSC (Fig. **5C**). Across the nMSC and CA-MSC lines used in the Cell Mito Stress Test, WT1 protein expression showed a positive correlation with basal respiration, maximal respiration, and proton leak (Fig. **S4A**), suggesting an association between WT1 expression and the CA-MSC mitochondrial phenotype.

To determine whether WT1 contributes functionally to this phenotype, we generated doxycycline-inducible WT1-overexpressing nMSC and CA-MSC with WT1 knockdown using short hairpin RNA (shRNA). WT1 overexpression was validated by western blotting (Fig.**5D**). Prolonged WT1 overexpression in nMSC significantly increased basal, maximal, and ATP-linked respiration, as well as proton leak (Fig. **5D-H**), accompanied by increased mitochondrial superoxide (Fig. **5N**). However, coupling efficiency remained unchanged (Fig. **S4E**), and WT1 overexpression increased rather than decreased mitochondrial membrane potential (Fig **S4D**). WT1 overexpression alone did not increase glycolytic activity in nMSC (Fig. **5P**). Together, these findings indicate that WT1 overexpression recapitulates several features of the CA-MSC mitochondrial phenotype, including increased respiratory activity, proton leak, and oxidative stress, but does not fully reproduce the mitochondrial state observed in established CA-MSC, or its glycolytic phenotype. Conversely, WT1 knockdown (KD) in CA-MSC, validated by western blotting (Fig. **5I**), reduced basal respiration, maximal respiration, and proton leak, while ATP-linked respiration remained unchanged (Fig. **5J-M**). WT1 KD also reduced mitochondrial superoxide (Fig. **5O**) as well as glycolytic activity (Fig. **5Q**), supporting a role for WT1 in maintaining both mitochondrial and glycolytic features of the established CA-MSC metabolic state.

## PDK4 contributes to the increased glycolytic activity of CA-MSC

To identify additional metabolic factors contributing to the CA-MSC phenotype, we examined differentially expressed metabolism-associated genes in our bulk RNA-seq dataset comparing nMSC and CA-MSC and identified PDK4 as a highly upregulated candidate (Fig. **S4F**). Increased PDK4 expression in CA-MSC was validated at the protein level by Western blotting and densitometric quantification (Fig. **5R**). PDK4 inactivates pyruvate dehydrogenase, blocking conversion of pyruvate to Aceytl CoA in the mitochondria, thereby limiting pyruvate oxidation, and conversely promoting fatty acid oxidation and a switch from glucose to lactic acid production in both cancer and stromal cells^34,35^. We therefore asked whether increased PDK4 expression could drive components of the CA-MSC metabolic phenotype. PDK4 overexpression in nMSC (Fig. **5U**) did not consistently alter mitochondrial respiratory parameters or mitochondrial ROS (Fig. **S4H–N**). However, as expected, PDK4 overexpression increased extracellular acidification rate (ECAR; Fig. **5V**), supporting a more specific contribution of PDK4 to the glycolytic component of CA-MSC metabolic reprogramming.

Given the reduction in glycolytic activity following WT1 knockdown, we next examined whether WT1 manipulation also affected PDK4 expression. WT1 knockdown in CA- MSC significantly reduced PDK4 protein expression (Fig. **5S**), whereas WT1 overexpression in nMSC did not significantly increase PDK4 expression (Fig. **5T**), suggesting that WT1 overexpression alone is insufficient to induce PDK4 expression. We next examined the functional consequences of PDK4 knockdown in CA-MSC. PDK4 knockdown in CA-MSC, confirmed by Western blotting (Fig. **5W**), significantly reduced cell proliferation compared with scramble controls (Fig. **5X**), preventing sufficient cell expansion for downstream metabolic assays. These findings suggest that elevated PDK4 expression may represent a compensatory metabolic adaptation to inefficient mitochondrial respiration and increased oxidative stress in CA-MSC, potentially supporting their growth and survival by favoring glycolytic metabolism.

## Oxidatively stressed CA-MSC mitochondria are preferentially transferred to ovarian cancer cells

We next asked whether the mitochondrial state/dysfunction influences the ability of mitochondria to be transferred from CA-MSC to ovarian cancer cells. We previously demonstrated that CA-MSC transfer mitochondria to ovarian cancer cells and that this transfer enhances cancer cell respiration, proliferation, chemoresistance, and metastatic potential^22^. We therefore asked whether mitochondria transferred from CA-MSC represent a randomly selected mitochondrial population or are enriched for the oxidatively stressed mitochondrial state.

Ratiometric imaging of CA-MSC expressing mitochondrial matrix-targeted HyPer7 (mitoHyper7) revealed a population of highly oxidized mitochondria, frequently localized toward the cell periphery (Fig. **6A**). Following coculture with ovarian cancer cells for 24 hours, highly oxidized CA-MSC derived mitochondria were detected within recipient tumor cells (Fig. **6B**).

**Figure 6.**
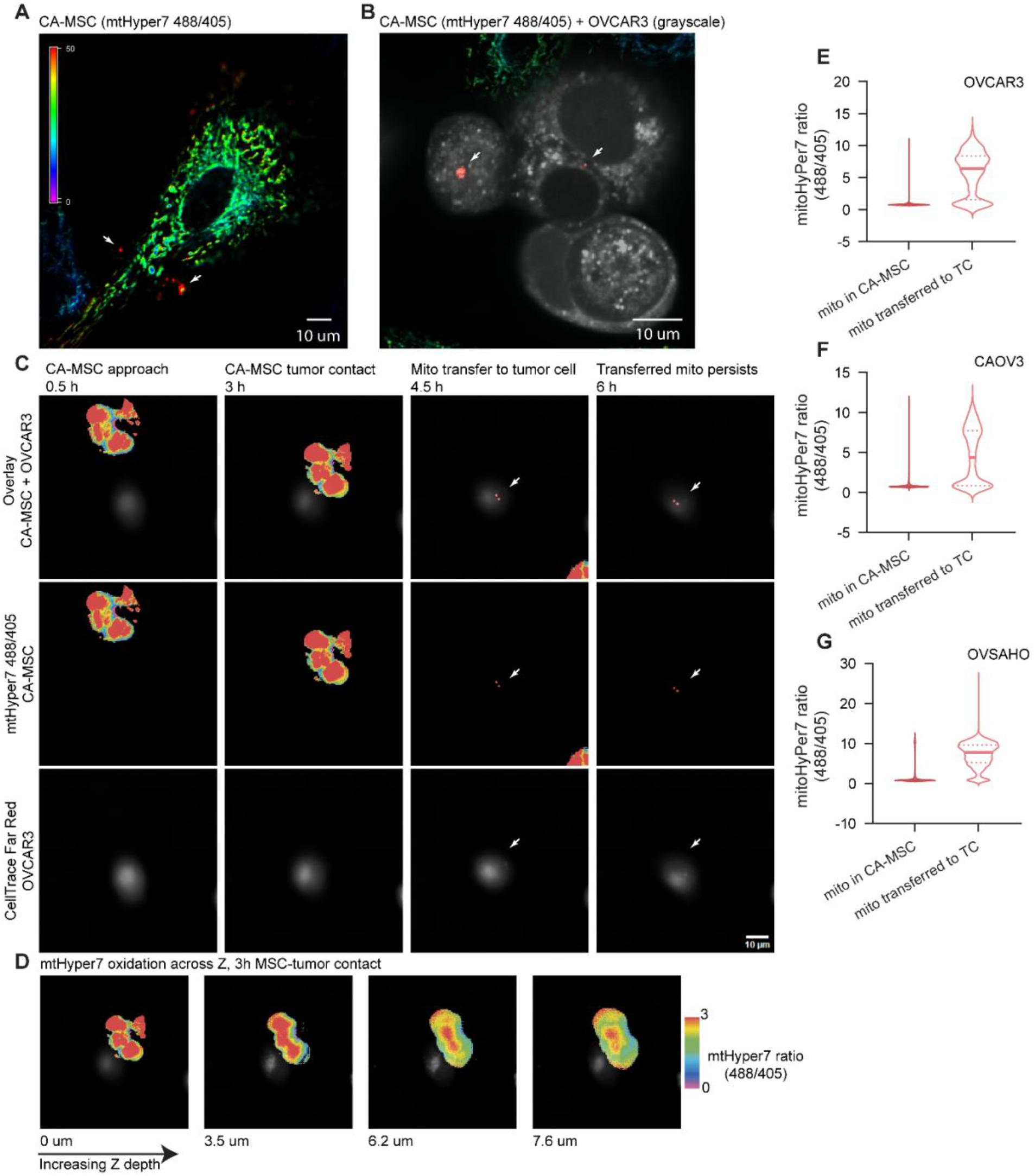
| CA-MSC transfer oxidatively stressed mitochondria to ovarian cancer cells. **A,B,** Representative ratiometric mitoHyPer7 images of CA-MSC cultured alone (A) and cocultured with Cell Trace Far Red (CTFR)-labeled OVCAR3 cells (B). HyPer7 ratio images were generated from the 488/405 nm signal, with red indicating the highest level of mitochondrial oxidation. CTFR-labeled tumor cells are shown in grayscale for visualization. White arrows indicate transferred CA-MSC mitochondria exhibiting elevated HyPer7 oxidation within recipient tumor cells. Scale bars, 10 μm. **C,** Representative time-lapse fluorescence images showing mitochondrial transfer from CA-MSC to CTFR-labeled OVCAR3 cells. Images capture CA-MSC approach (0.5 h), contact with the tumor cell (3 h), mitochondrial transfer (4.5 h), and persistence of the transferred mitochondrial signal (8 h). Merged images (top) show CA-MSC mitoHyPer7 and CTFR-labeled OVCAR3 cells, with individual mitoHyPer7 (middle) and CTFR (bottom) channels shown separately. White arrows indicate transferred CA-MSC mitochondria. Scale bar, 10 μm. **D,** Representative ratiometric mitoHyPer7 images acquired at increasing Z-depths (0, 3.5, 6.2, and 7.6 μm) during CA-MSC–OVCAR3 contact at 3 h, showing the mitochondrial oxidation signal across different optical sections. Red indicates the highest HyPer7 oxidation ratio. **E–G,** Flow cytometric analysis of mitochondrial HyPer7 oxidation ratios in mitochondria retained within CA- MSC and CA-MSC mitochondria following transfer into recipient OVCAR3 (E), CAOV3 (F), and OVSAHO (G) ovarian cancer cells. Violin plots show the distribution of single- cell HyPer7 ratios (488/405) from a representative coculture experiment for each tumor cell line. HyPer7 ratios in (E–G) were calculated from exported single-cell flow cytometry data. Quantification of geometric mean HyPer7 ratios from three independent coculture experiments for each tumor cell line is provided in Supplementary Fig. 4O.

To determine whether CA-MSC mitochondria were already oxidized at the time of transfer rather than becoming oxidized after entering tumor cells, we performed live-cell time-lapse imaging of CA-MSC–tumor cell interactions (**Supplementary Video 1**).

Time-lapse imaging captured CA-MSCs with oxidized mitochondria approaching and contacting tumor cells, followed by transfer of the oxidized CA-MSC mitochondria to the tumor cell, which remained detectable after transfer (Fig. **6C**; **Supplementary Video 1**). Examination of different optical Z-sections of the same donor CA-MSC during the CA-MSC-tumor contact (3 h) revealed heterogeneity in mitochondrial oxidation (Fig. **6D**) indicating that there was selective transfer of oxidized CA-MSC mitochondria to tumor cells.

To quantify the oxidative state of transferred mitochondria, we performed coculture under ULA conditions, where mitochondrial transfer occurs robustly, and measured mitoHyPer7 ratios by flow cytometry. Of note, only CA-MSC were transduced with mitoHyPer7 thus only CA-MSC derived mitochondria carried this fluorescent label and were quantified. Across co-culture with three HGSOC tumor cell lines (OVCAR3, CAOV3, OVSAHO), CA-MSC mitochondria detected within recipient tumor cells exhibited significantly higher mitoHyPer7 ratios than mitochondria retained within donor CA-MSC (Fig. **6E–G****, S4O**). Together, these findings support the preferential transfer of oxidatively stressed CA-MSC mitochondria to ovarian cancer cells.

## Mitochondrial variant tracking identifies a tumor subpopulation with stroma- associated mitochondria and a distinct transcriptional state

To determine whether the stromal-tumor mitochondrial donation observed in our experimental models could also be detected within patient tumors, we performed single- cell RNA sequencing (scRNAseq) of a freshly dissociated ovarian tumor (directly processed into single cells without culture). CA-MSC and tumor cells were also isolated from the same tumor, briefly cultured and validated for cell type identity. These cells were used to inform cell identity in the scRNAseq analysis but not used for down-stream analysis (Fig. **S5A**).

Focusing on the tumor and stromal cell compartments from the bulk tumor, clustering of the scRNAseq dataset identified multiple subpopulations within both tumor and stromal populations (Fig. **7A**). Tumor cells separated into five clusters (2, 6, 7, 10, and 11), whereas stromal cells separated into three clusters (4, 5, and 9). To further assess the distinction between malignant and nonmalignant cell populations, we performed inferCNV analysis using immune cell populations as references to determine genomic copy-number variations. The analysis revealed copy-number alterations in the annotated tumor cells relative to the reference populations, supporting their malignant identity (Fig. **S5B**) and lack of copy-number variants in the stromal populations supporting their non-malignant identity. Mitochondrial variant tracking revealed that tumor Cluster 6 shared greater similarity in mitochondrial variant patterns with stromal clusters 4, 5, and 9 than with the other tumor populations (Fig. **7B**). These findings identify a tumor subpopulation within the patient tumor with mitochondrial features closely related to those of stromal cells supporting the relevance of in vivo stromal to tumor cell mitochondrial transfer. Notably, Cluster 6 accounted for approximately 21% of the annotated tumor cells in this specimen, indicating that the mitochondrially distinct subpopulation represented a substantial proportion of the tumor cells analyzed.

**Figure 7.**
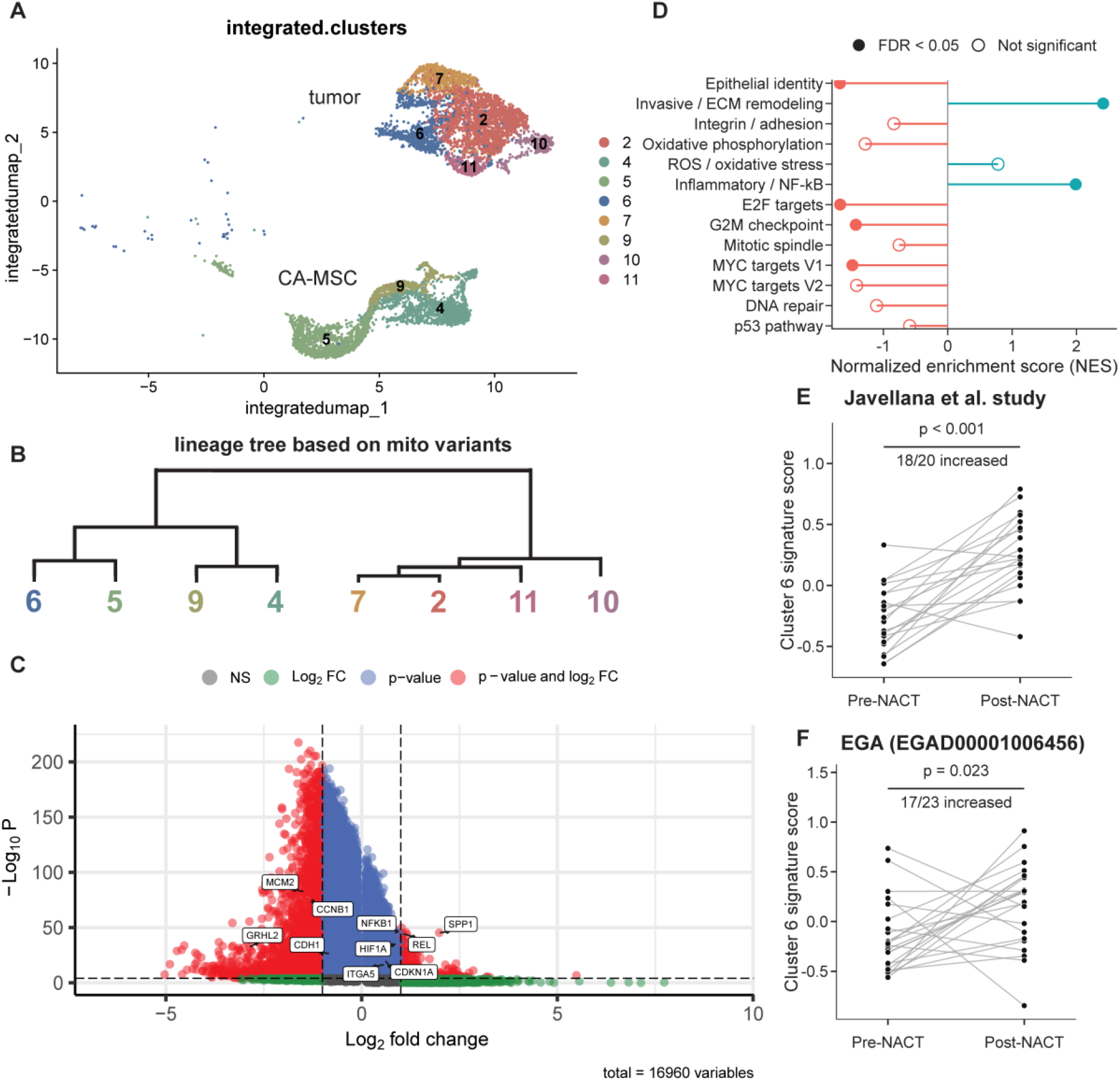
| A distinct HGSOC tumor subpopulation shares mitochondrial variant patterns with CA-MSC and exhibits a transcriptional signature enriched in post- chemotherapy residual tumors. **A,** Uniform manifold approximation and projection (UMAP) representation of single-cell RNA sequencing (scRNA-seq) data from a patient-derived HGSOC tumor, showing tumor and carcinoma-associated mesenchymal stem cell (CA-MSC) populations. Tumor Cluster 6 is highlighted by a red circle. **B,** Mitochondrial variant-based lineage tree reconstructed from the same scRNA-seq dataset, showing relationships among tumor and CA-MSC clusters. **C,** Volcano plot showing differential gene expression between tumor Cluster 6 and tumor Clusters 2, 7, 10, and 11. **D,** Preranked gene set enrichment analysis (GSEA) of 10 Molecular signatures database (MSigDB) Hallmark pathways and 3 custom gene modules (epithelial identity, invasive/extracellular matrix (ECM) remodeling, and integrin/adhesion), using the full Cluster 6 versus other tumor-cluster differential expression results ranked by average log₂ fold change. Positive normalized enrichment scores (NES) indicate enrichment in Cluster 6, with filled circles indicating statistically significant pathways (FDR < 0.05). **E,F,** Comparison of Cluster 6-like transcriptional signature scores between patient-matched pre-neoadjuvant chemotherapy (pre-NACT) tumors and post-NACT residual tumors from the Javellana HGSOC cohort (E; n = 20 pairs) and EGA HGSOC cohort (F; n = 23 pairs). Signature scores were calculated using both upregulated and downregulated genes identified in Cluster 6. For A–D, analyses were performed using scRNA-seq data from a single patient-derived HGSOC tumor. Differential gene expression between tumor Cluster 6 and the remaining tumor clusters (2, 7, 10, and 11) was assessed using the Seurat FindMarkers function with a Wilcoxon rank-sum test. Genes with an adjusted P < 0.05 and an absolute average log₂ fold change ≥ 0.5 were considered differentially expressed. For E,F, 20 matched pre- and post-NACT tumor pairs were analyzed in the Javellana cohort (E), with increased Cluster 6-like signature scores observed in 18 of 20 pairs (*P* < 0.001), and 23 matched pairs were analyzed in the EGA cohort (F), with increased signature scores observed in 17 of 23 pairs (*P* = 0.023). Statistical significance was determined using two-sided paired Wilcoxon signed-rank tests.

We next asked whether this tumor population also exhibited a distinct transcriptional state. Differential expression analysis comparing tumor Cluster 6 with the remaining tumor clusters (2, 7, 10, and 11) revealed increased expression of genes associated with inflammatory signaling (REL, NF-kB1), adhesion and microenvironmental interactions (ITGA5, SPP1), and cellular stress responses (HIF1A). In contrast, Cluster 6 exhibited reduced expression of cell-cycle-associated genes (MCM2, CCNB1) and epithelial identity markers (GRHL2, CDH1) (Fig. **7C**). Pathway enrichment analysis further revealed distinct biological programs associated with Cluster 6. Kyoto Encyclopedia of Genes and Genomes (KEGG) analysis identified pathways related to immune responses, antigen processing and presentation, and cell adhesion molecules among upregulated genes, whereas downregulated genes were associated with cell- cycle regulation, DNA replication, and cadherin signaling (Fig. **S5C,D**). Consistent with these findings, gene set enrichment analysis (GSEA) revealed enrichment of invasive and extracellular matrix remodeling and inflammatory/NF-κB-associated programs, together with depletion of epithelial identity and cell-cycle-associated programs, including E2F targets and G2M checkpoint (Fig. **7D**). Together, these findings indicate that tumor Cluster 6 exhibits a distinct transcriptional state characterized by altered epithelial identity, cell adhesion and extracellular matrix interactions, inflammatory signaling, and reduced cell-cycle-associated programs.

Given this distinct transcriptional profile, we next asked whether a similar state could be identified in independent HGSOC patient cohorts, particularly in residual tumors following chemotherapy, which are enriched in therapy-resistant cells. We generated a Cluster 6 transcriptional signature using genes significantly upregulated and downregulated in Cluster 6 relative to the other tumor clusters and examined its expression in four independent cohorts of patient-matched tumors collected before and after neoadjuvant chemotherapy (NACT)^36,37,38–40^. In the Javellana cohort, 18 of 20 patients showed higher Cluster 6-like signature scores following NACT (paired Wilcoxon, *P* < 0.001; Fig. **7E**). A similar increase was observed in the EGA cohort, where 17 of 23 patients exhibited increased signature scores in post-NACT residual tumors compared with matched pre-NACT samples (paired Wilcoxon, *P* = 0.023; Fig. **7F**). These findings were further supported by two additional independent cohorts, GSE143897 and GSE227100, where 14 of 18 patients (*P* = 0.0407; Supplementary Fig. **5E**) and 17 of 24 patients (*P* = 0.0268; Supplementary Fig. **5F**), respectively, exhibited increased Cluster 6-like signature scores following NACT.

Together, these findings identify a transcriptionally distinct tumor subpopulation sharing mitochondrial variant features with the stromal/CA-MSC compartment. The enrichment of a similar transcriptional signature in post-NACT tumors suggests that this cellular state may also be relevant to residual ovarian cancer following chemotherapy and the development of therapy resistance.

## Discussion

Our findings define a distinct metabolic state in CA-MSC characterized by increased bioenergetic activity despite multiple features of mitochondrial stress. Rather than exhibiting the severe bioenergetic failure associated with overt mitochondrial dysfunction, CA-MSC maintain or increase respiratory capacity while operating at reduced energetic efficiency. Increased glycolytic activity may provide an additional means of meeting cellular energy demands despite inefficient mitochondrial respiration. This simultaneous increase in oxidative and glycolytic metabolism therefore suggests broad metabolic adaptation rather than a simple shift between energetic pathways. The coexistence of elongated and more extensively branched mitochondria with increased punctate mitochondrial structures, greater total mitochondrial area, and increased mitochondrial number further suggests heterogeneity within the CA-MSC mitochondrial population. CA-MSC may therefore accumulate stressed or functionally compromised mitochondria alongside mitochondria capable of sustaining or potentially increasing respiratory activity to compensate, allowing these cells to maintain high metabolic activity despite increased oxidative stress and reduced mitochondrial efficiency.

Mitochondrial genomic alterations may represent an additional component of this stress- adapted mitochondrial state whether it is the cause or the consequence. CA-MSC exhibited an increased mtDNA variant burden together with enrichment of homoplasmic variants, which may have greater potential to influence mitochondrial function. The relationship between these genomic alterations and the mitochondrial phenotype, however, remains unclear. As seen in literature, chronic mitochondrial oxidative stress could contribute to mtDNA damage, although whether this contributes to the increased mtDNA variant burden observed in CA-MSC remains unclear^41^. Conversely, functionally relevant mtDNA variants have been shown to alter mitochondrial function and increase oxidative stress, potentially contributing to the mitochondrial phenotype observed in CA- MSC^42^. Mitochondrial genomic alterations and oxidative stress may therefore represent interconnected features of CA-MSC reprogramming, although the relationship between these processes remains to be determined. While CA-MSC exhibited an overall increase in mtDNA variant burden and enrichment of homoplasmic variants, the number and distribution of variants varied among individual patient-derived MSC lines in both bulk and single-cell datasets. The relatively small number of samples in the single-cell analysis (two nMSC and three CA-MSC) limits our ability to determine whether specific mtDNA variants reproducibly characterize CA-MSC across patients. Larger patient cohorts will be needed to determine whether any recurrent variants or mutational patterns are associated with the CA-MSC state.

Because individual mtDNA variants cannot be readily introduced into nMSC to directly determine their functional consequences, we examined ETC function in the same patient-derived MSC lines used for our single-cell ATAC-seq analysis. The complex- specific respiratory and transcriptional alterations observed in CA-MSC suggest that these cells actively maintain oxidative metabolism despite mitochondrial stress.

Seahorse analysis of intact MSC revealed increased maximal respiration in CA-MSC despite reduced coupling efficiency, whereas analysis of isolated mitochondria using equal mitochondrial input revealed largely maintained Complex I/II-supported respiration with increased proton leak under Complex I-supported conditions, but reduced downstream Complex III/IV-supported activity. Notably, the CA-MSC line harbored a likely pathogenic variant in MT-ND1 (3715G>A), which encodes a core subunit of Complex I involved in proton translocation. The increased proton leak observed under Complex I-supported conditions raises the possibility that this variant contributes to altered respiratory coupling, although its direct functional consequences remain to be established. Increased mitochondrial abundance may therefore contribute to the higher respiratory activity observed at the whole-cell level. Reduced downstream respiratory capacity also raises the possibility that Complex III/IV function represents a bottleneck within the CA-MSC ETC, while maintenance of upstream respiration may reflect compensatory adaptation to preserve respiratory activity. Consistent with this model, transcriptional enrichment of OXPHOS and TCA cycle programs further supports coordinated upregulation of oxidative metabolism in CA-MSC.

Increased respiratory activity may also contribute to mitochondrial stress over time. Previous studies have demonstrated that elevated mitochondrial metabolic activity can be accompanied by increased ROS production and longer-term cellular stress in metabolically active stromal and other cell populations^3,43,44^. In CA-MSC, sustained respiratory activity in the setting of altered ETC function may similarly contribute to increased oxidative stress and proton leak. ROS-dependent stimulation of mitochondrial proton conductance could further reinforce this phenotype^27,28^. Thus, increased oxidative metabolism may represent both a compensatory response that enables CA- MSC to maintain respiratory activity and a potential contributor to mitochondrial stress. Whether increased respiration precedes oxidative stress or develops as a compensatory response remains unresolved, and these processes may reinforce one another during CA-MSC reprogramming.

The increased mitochondrial abundance observed in CA-MSC may have additional significance given their role as mitochondrial donors^22^. CA-MSC must maintain their own bioenergetic requirements while also transferring mitochondria to ovarian cancer cells^22^. Expansion of the mitochondrial compartment could therefore provide a larger mitochondrial pool from which mitochondria can be transferred without compromising stromal cell bioenergetic capacity. Whether increased mitochondrial abundance specifically supports mitochondrial transfer or instead reflects a broader adaptation to increased metabolic demand remains to be determined.

Our findings identify WT1 as an important regulator of the CA-MSC mitochondrial state. Prolonged WT1 overexpression increased mitochondrial respiration, proton leak, and oxidative stress in nMSC, recapitulating several central features of the CA-MSC mitochondrial phenotype. The time required for these changes to emerge is consistent with the progressive nature of CA-MSC reprogramming, as nMSC exposed to ovarian cancer cells similarly require several days to acquire CA-MSC-like properties^17,21^. This temporal requirement may reflect sustained transcriptional reprogramming and/or the gradual development of mitochondrial stress. However, WT1 overexpression did not reproduce all features of established CA-MSC: coupling efficiency remained unchanged and mitochondrial membrane potential increased rather than decreased showing the partial depolarization observed in CA-MSC. Conversely, WT1 depletion disrupted the mitochondrial and glycolytic features of the established CA-MSC state. Together, these findings support WT1 as a major driver of mitochondrial reprogramming while suggesting that additional factors or prolonged tumor-associated stress are required to establish the complete CA-MSC metabolic phenotype.

The relationship between WT1 and PDK4 also appears to be indirect. Although WT1 depletion reduced PDK4 expression, WT1 overexpression was not sufficient to induce PDK4, arguing against a simple model of direct WT1-dependent regulation. Instead, PDK4 induction may emerge as a secondary or compensatory response to the mitochondrial and metabolic changes that develop during CA-MSC reprogramming. In this model, reversal of the mitochondrial phenotype following WT1 depletion would also remove the cellular state that sustains elevated PDK4 expression. This interpretation is consistent with PDK4 contributing to the glycolytic phenotype of CA-MSC without being directly induced by WT1. Notably, PDK4 knockdown significantly impaired CA-MSC proliferation, preventing sufficient cell expansion for downstream metabolic assays.

These findings indicate that elevated PDK4 represents an important compensatory adaptation necessary to sustain CA-MSC growth and survival in the context of their altered mitochondrial metabolism.

Importantly, the progressive increase in mitochondrial oxidation under tumor-associated stress conditions indicates that this mitochondrial phenotype is not simply an intrinsic property of CA-MSC but can develop as nMSC become reprogrammed into CA-MSCs in response to the ovarian tumor microenvironment. Tumor cell exposure alone was sufficient to increase mitochondrial oxidation, whereas anchorage-independent, nutrient-deprived conditions also induced oxidation even in the absence of tumor cells. Combining tumor exposure with these environmental stresses produced the strongest oxidative phenotype, suggesting that tumor-derived signals and microenvironmental stress cooperate to establish the CA-MSC mitochondrial state. This may be particularly relevant during ovarian cancer dissemination, where tumor cells and stromal cells exist as multicellular aggregates within ascites and encounter anchorage-independent and metabolically challenging conditions^21,45,46^. Although ULA and serum deprivation do not fully recapitulate this environment, the acquisition of a stress-adapted mitochondrial state under these conditions raises the possibility that stromal mitochondrial reprogramming facilitates adaptation to periods of increased microenvironmental stress.

The preferential transfer of oxidatively stressed mitochondria suggests that the CA-MSC mitochondrial phenotype may have consequences beyond the stromal population. We previously demonstrated that mitochondrial transfer from CA-MSC increases ovarian cancer cell respiration, proliferation, chemoresistance, metastatic capacity, and tumor cell heterogeneity^22^. In the current study, transferred CA-MSC mitochondria exhibited a higher oxidative state than mitochondria retained within donor CA-MSC. Importantly, live-cell time-lapse imaging revealed that mitochondria were already highly oxidized during transfer, rather than becoming oxidized after entering the tumor cell. Examination of different optical sections also revealed heterogeneity in mitochondrial oxidation within different planes in the same donor CA-MSC, supporting the idea of preferential transfer of an oxidized mitochondrial population. These findings raise the possibility that CA- MSC provide tumor cells not only with additional mitochondrial bioenergetic capacity, but also with mitochondria carrying a distinct redox state. Given the role of mitochondrial ROS in regulating cellular stress responses and metabolic adaptation^9,10^, the transfer of oxidized mitochondria could potentially influence the adaptive state of recipient tumor cells. However, whether the oxidative state of transferred mitochondria directly contributes to the tumor-promoting effects of mitochondrial transfer remains to be determined.

Importantly, our mitochondrial variant tracking analysis of freshly derived HGSOC tissue provides further insight into the physiologic relevance of tumor–stromal mitochondrial exchange in patients. We identified a tumor subpopulation sharing mitochondrial variant patterns with the stromal population, consistent with the stromal to tumor mitochondrial exchange observed in our experimental models. Interestingly, this mitochondrially related tumor population, identified as Cluster 6, exhibited a distinct transcriptional state compared with other tumor populations. Differential expression and pathway analyses revealed increased expression of genes associated with inflammatory NF-kB signaling, and invasive/extracellular matrix remodeling, alongside reduced expression of epithelial identity and cell-cycle-associated pathways. These transcriptional differences suggest that Cluster 6 may represent a tumor cell state with altered interactions with its microenvironment and adaptations to cellular stress. Notably, Cluster 6 represented approximately 21% of the annotated tumor cell population in the patient tumor analyzed, indicating that this mitochondrially distinct population was a not a rare subset within this tumor. The presence of a substantial tumor population sharing mitochondrial features with the stromal compartment suggests that tumor–stromal mitochondrial relationships are relevant to intratumoral heterogeneity and may drive specific tumor programs.

The clinical relevance of this transcriptional state was further supported by its enrichment in residual tumors following neoadjuvant chemotherapy. Across four independent patient-matched HGSOC cohorts, post-NACT tumors exhibited increased expression of the Cluster 6 derived transcriptional signature compared with matched pre-NACT tumors. Given the reduced expression of cell-cycle-associated genes and increased expression of stress- and inflammatory signaling-associated genes in Cluster 6, this finding raises the possibility that similar transcriptional programs may be associated with tumor persistence/resistance to chemotherapy and may be important drivers in ovarian cancer recurrence and overall survival^47,48^. However, the enrichment of this signature in bulk tumors does not establish whether chemotherapy selectively enriches a pre-existing tumor population or induces transcriptional changes in surviving cells. Furthermore, whether mitochondrial transfer contributes to the emergence or maintenance of this transcriptional state remains to be determined, although we have previously shown that mitochondrial transfer from CA-MSC to ovarian cancer cell leads to intratumoral heterogeneity and therapy resistance^22^. Together, these findings suggest that CA-MSC mitochondrial reprogramming and the transfer of oxidatively stressed mitochondria contribute to tumor mitochondrial heterogeneity and influence distinct tumor cell states within the ovarian cancer microenvironment.

## Methods

### nMSC and CA-MSC isolation and culture

Primary human normal mesenchymal stromal/stem cells (nMSC) and cancer-associated mesenchymal stromal/stem cells (CA-MSC) were isolated and characterized as previously described^15,21,49^. Human tissue samples were obtained under protocols approved by the University of Pittsburgh Institutional Review Board (STUDY22070111), with informed consent obtained from all participants. nMSC were derived from gynecologic surgical specimens from patients undergoing surgery for benign indications, whereas CA-MSC were isolated from surgical specimens from patients with high-grade serous ovarian cancer (HGSOC).

Fresh tissue was minced into <1 mm³ fragments and plated in six-well tissue culture plates. In accordance with the International Society for Cell and Gene Therapy criteria for defining multipotent mesenchymal stromal cells, MSCs were selected based on plastic adherence and expression of the cell-surface markers CD105, CD90, and CD73. Adipogenic, osteogenic, and chondrogenic differentiation capacity was verified as previously described^16,17,21.^

MSCs were maintained in Mammary Epithelial Cell Basal Medium (MEBM) supplemented with 10% heat-inactivated fetal bovine serum (HI-FBS), 1× B27 supplement, 20 ng/mL epidermal growth factor, 1 ng/mL hydrocortisone, 5 μg/mL insulin, 100 μM β-mercaptoethanol, 10 ng/mL basic fibroblast growth factor, 1% penicillin–streptomycin, and 20 μg/mL gentamicin. Cells were maintained at 37 °C in a humidified incubator with 5% CO₂ and were used for experiments between passages [P2–P8].

## Ovarian cancer cell lines and culture

Human ovarian cancer cell lines OVCAR3, CAOV3, and OVSAHO were used in this study. OVCAR3 cells were maintained in RPMI 1640 supplemented with 20% HI-FBS, 1% penicillin–streptomycin, and 1:10,000 bovine insulin, and CAOV3 cells were maintained in RPMI 1640 supplemented with 10% HI-FBS and 1% penicillin– streptomycin, and OVSAHO were maintained in Dulbecco’s eagle modified medium (DMEM) supplemented with 10% HI-FBS and 1% penicillin–streptomycin. All cells were cultured at 37 °C in a humidified incubator with 5% CO₂.

Cell lines were routinely tested for mycoplasma contamination and were used only when negative. Cell-line identity was confirmed by short tandem repeat (STR) profiling. For coculture experiments, ovarian cancer cells were fluorescently labeled as described below to distinguish tumor cells from MSC populations.

## Lentiviral transduction and generation of WT1- and PDK4-modified MSCs

To assess the functional contribution of WT1 and PDK4 to the CA-MSC metabolic phenotype, WT1 and PDK4 were depleted in CA-MSC by short hairpin RNA (shRNA)- mediated knockdown and overexpressed in nMSC using lentiviral constructs. For knockdown experiments, CA-MSC were transduced with lentiviral particles encoding shRNAs targeting WT1 or PDK4. A non-targeting scrambled shRNA was used as the corresponding control.

For overexpression studies, nMSC were transduced with lentiviral constructs encoding WT1-A or PDK4, with the corresponding empty vectors used as controls. The WT1-A construct was modified by GenScript to incorporate a doxycycline-inducible expression backbone, permitting temporal control of WT1 expression. Following establishment of stable transduced cells, WT1 expression was induced with 100ng/ml doxycycline for at least 5 days before experimental analysis. Control cells transduced with the corresponding control vector were treated with the same concentration and duration of doxycycline.

For lentiviral transduction, MSCs were seeded at 25,000 cells per well in six-well tissue culture plates and transduced at a multiplicity of infection (MOI) ranging from 10 to 50. Viral medium was removed 24 h after transduction and replaced with complete MSC medium. Transduced cells were subsequently selected with 0.4 μg/mL puromycin for 3 days. Following selection, cells were allowed to recover in complete MSC medium before downstream experimental analysis. WT1 and PDK4 knockdown or overexpression was confirmed by immunoblotting before downstream analyses.

To assess the effect of PDK4 knockdown on CA-MSC proliferation, equal numbers of scramble control and PDK4-knockdown CA-MSC were seeded under identical culture conditions. Cells were harvested 48 h after seeding and counted using trypan blue exclusion and a hemocytometer. Cell proliferation was expressed as fold change in cell number relative to the initial seeding density.

## Seahorse extracellular flux analysis

Cellular bioenergetics were assessed using an Agilent Seahorse XF HS Mini Extracellular Flux Analyzer (Agilent Technologies). To minimize differences in cell number arising from variable proliferation rates among primary MSC lines, cells were collected on the day of the assay, counted, and resuspended directly in the corresponding Seahorse assay medium. An equal number of cells (5,000 cells per well) was seeded in 80 μL of assay medium into poly-D-lysine (PDL)-coated Seahorse XF HS Mini 8-well miniplates. Plates were centrifuged at 800 rpm for 2 min to promote uniform attachment of cells to the bottom of the wells and formation of a monolayer. Cells were then incubated at 37 °C in a humidified non CO_2_ incubator for 45 min. Immediately before initiation of the assay, an additional 100 μL of the corresponding Seahorse assay medium was gently added to each well, bringing the final assay volume to 180 μL.

For the Cell Mito Stress Test, cells were assayed in Seahorse XF DMEM assay medium supplemented with glucose (10 mM), glutamine (2 mM), and sodium pyruvate (1 mM). Oxygen consumption rate (OCR) and extracellular acidification rate (ECAR) were measured at baseline and following sequential injection of oligomycin (1.5 μM), FCCP (2.5 μM), and antimycin A/rotenone (0.5 μM each). Basal respiration, ATP-linked respiration, proton leak, maximal respiration, spare respiratory capacity, and coupling efficiency were calculated from the resulting OCR measurements according to the manufacturer’s Cell Mito Stress Test protocol.

For the Glycolysis Stress Test, cells were assayed in Seahorse XF DMEM assay medium supplemented with glutamine (2 mM) in the absence of glucose and pyruvate. ECAR was measured following sequential injection of glucose (10 mM), oligomycin (1 μM), and 2-deoxy-D-glucose (2-DG; 50 mM). Glycolysis was calculated according to the manufacturer’s Glycolysis Stress Test protocol.

Because post-assay normalization was not feasible for these primary cell lines, extracellular flux measurements were compared between groups using an equal number of cells seeded per well at the start of each assay.

## Confocal imaging and analysis of mitochondrial morphology

To visualize mitochondrial morphology, nMSC and CA-MSC were transduced with a lentiviral construct encoding mGFP fused to the mitochondrial targeting sequence of cytochrome c oxidase subunit VIII (COX8-GFP) at a multiplicity of infection (MOI) of 10. Following transduction, cells were seeded into μ-Slide 4 well (Ibidi) and allowed to adhere overnight. The following day, cells were fixed with a solution containing 4% paraformaldehyde and 2.5% glutaraldehyde in phosphate-buffered saline (PBS) for 20 min at room temperature. Cells were subsequently washed three times with PBS and imaged by confocal microscopy using a 63× oil-immersion objective (Nikon AX Confocal).

Three independent patient-derived nMSC lines and three independent patient-derived CA-MSC lines were analyzed. Z-stacks encompassing the mitochondrial network were acquired and converted to two-dimensional maximum-intensity projections for quantitative analysis.

Mitochondrial network morphology was analyzed in ImageJ/Fiji using the Mitochondria Analyzer plugin^50^ with optimized two-dimensional (2D) thresholding parameters.

Maximum-intensity projections of COX8-GFP fluorescence images were processed using a custom ImageJ macro to automate image preprocessing and thresholding. Briefly, images were converted to 8-bit grayscale, contrast was enhanced (0.35% saturated pixels), and a sharpening filter was applied. Background subtraction was performed using a rolling-ball radius of 1.25 μm, followed by Sigma Filter Plus (radius 0.71 pixels), local contrast enhancement (maximum slope 1.8), and gamma adjustment (0.8). Adaptive thresholding was performed using the Weighted Mean method with a block size of 2.5 μm and a C-value of 10, followed by despeckling and outlier removal (radius 0.4 pixels, threshold 50). The same optimized thresholding parameters were applied consistently across all nMSC and CA-MSC images.

Following thresholding, individual cells were manually outlined using regions of interest (ROIs), and mitochondrial morphology was quantified using the Mitochondria Analyzer 2D Analysis function, with a minimum mitochondrial object area of 0.06 μm².

Morphological parameters included mitochondrial object count, total mitochondrial area, mean aspect ratio, mean form factor, branch length per mitochondrial object, and number of branches per mitochondrial object.

Punctate mitochondrial structures were quantified separately from the original 16-bit COX8-GFP maximum-intensity projection images using a custom image-analysis workflow. The same manually defined cell ROIs were used to identify bright punctate structures based on fluorescence intensity, size, shape, and surrounding intensity profiles. Candidate puncta were identified as local fluorescence intensity maxima and classified using a minimum core area of 0.158 μm², aspect ratio ≤ 1.7, circularity ≥ 0.85, solidity ≥ 0.90, and a compact, locally isotropic intensity profile. These parameters were optimized using manually annotated punctate structures and subsequently applied consistently across all nMSC and CA-MSC images without using cell-line identity during classification. Elongated tubular structures, branch points, and uncertain structures were excluded, and the number of punctate mitochondrial structures per cell was quantified.

A total of 186 cells from three independent patient-derived nMSC lines and three independent CA-MSC lines were included in the analysis. Mitochondrial morphology and punctate mitochondrial structures were compared between one nMSC and one CA- MSC line, with two additional independent nMSC and CA-MSC line comparisons performed to assess the consistency of the findings. Statistical analyses were performed using two-sided, unpaired Student’s *t*-tests, with individual cells treated as observations.

## Transmission electron microscopy

Transmission electron microscopy (TEM) sample preparation was performed at the University of Pittsburgh Center for Biological Imaging (CBI). Cell monolayers were briefly rinsed with 1× phosphate-buffered saline (PBS) and fixed in situ with 2.5% glutaraldehyde in 1× PBS (pH 7.4) for 1 h at room temperature. Samples were washed three times with PBS for 10 min each and post-fixed in 1% osmium tetroxide containing 1% potassium ferricyanide for 1 h at 4 °C. Following three additional PBS washes (10 min each), samples were dehydrated through a graded ethanol series (30%, 50%, 70%, and 90%; 10 min each), followed by three changes of 100% ethanol (15 min each).

Samples were subsequently infiltrated with Epon resin through three changes (1 h each). Resin-filled embedding capsules were inverted over the monolayers and polymerized overnight at 37 °C, followed by 48 h at 60 °C.

Semi-thin sections (300 nm) were prepared using a Reichert Ultracut ultramicrotome (Leica Microsystems), stained with 0.5% Toluidine Blue O in 1% sodium borate, and examined by light microscopy to identify regions of interest. Ultrathin sections (65 nm) were subsequently prepared, stained with 2% uranyl acetate (Electron Microscopy Sciences) and Reynolds’ lead citrate, and examined using a JEOL 1400 Plus transmission electron microscope equipped with a side-mounted AMT 2K digital camera (Advanced Microscopy Techniques).

Mitochondrial ultrastructure was assessed qualitatively in three independent patient- derived nMSC lines and three independent patient-derived CA-MSC lines. At least eight cells per patient line were imaged, and representative images were selected to illustrate mitochondrial ultrastructural features.

## Measurement of mitochondrial superoxide

Mitochondrial superoxide levels were measured using MitoSOX™ Red mitochondrial superoxide indicator. A standard 50 μg vial of lyophilized MitoSOX Red was reconstituted in 13 μL dimethyl sulfoxide (DMSO) to generate a 5 mM stock solution and subsequently diluted 1:1000 in Hank’s Balanced Salt Solution (HBSS) to a final working concentration of 5 μM.

Cells were trypsinized, pelleted and resuspended in HBSS with 5uM MitoSOX Red and incubated for 30 min at 37 °C in a non-CO₂ incubator. As a positive control for mitochondrial superoxide production, 50 μM antimycin A was added concurrently with MitoSOX Red during the staining period. Following incubation, cells were centrifuged, washed once with HBSS, and resuspended in flow cytometry buffer consisting of 2% fetal bovine serum in phosphate-buffered saline. Samples were maintained on ice and analyzed immediately by flow cytometry.

MitoSOX Red fluorescence was detected in the ECD-A channel using a Beckman Coulter CytoFLEX 4L flow cytometer. Flow cytometry data were analyzed using FlowJo version 10, and mitochondrial superoxide levels were quantified as the geometric mean fluorescence intensity (MFI) of the MitoSOX Red signal.

## Measurement of mitochondrial matrix oxidation using mitoHyPer7

Mitochondrial matrix hydrogen peroxide levels were measured using the genetically encoded ratiometric redox sensor Hyper7 targeted to the mitochondrial matrix (mitoHyPer7). nMSC and CA-MSC were transduced with lentiviral particles encoding mitoHyPer7 and expanded prior to analysis.

For comparison of basal mitochondrial oxidation between nMSC and CA-MSC, cells were seeded at 25,000 cells per well in six-well tissue culture plates. Culture medium was replaced the following day, and mitoHyPer7 fluorescence was measured by flow cytometry 24 h later.

mitoHyPer7 fluorescence was measured using excitation at 488 nm and 405 nm, with signals collected in the FITC and KO525 channels, respectively, on a Beckman Coulter CytoFLEX 4L flow cytometer. Flow cytometry data were analyzed using FlowJo version 10. For bulk analysis, mitochondrial oxidation was quantified as the ratio of the geometric mean fluorescence intensity obtained following 488-nm excitation to that obtained following 405-nm excitation (488/405 ratio). Higher 488/405 ratios were interpreted as increased oxidation of the mitoHyPer7 sensor.

## Measurement of mitochondrial membrane potential

Mitochondrial membrane potential was assessed using the ratiometric fluorescent dye JC-1. Cells were detached by trypsinization and resuspended in complete MSC culture medium containing JC-1 at a final concentration of 5 μg/mL. Cells were incubated for 30 min at 37 °C in a humidified 5% CO₂ tissue culture incubator. As a positive control for mitochondrial depolarization, cells were treated with 50 μM FCCP concurrently with JC- 1 staining.

Following staining, cells were washed once with phosphate-buffered saline and resuspended in flow cytometry buffer consisting of 2% fetal bovine serum in phosphate- buffered saline. Samples were maintained on ice and analyzed immediately by flow cytometry using a Beckman Coulter CytoFLEX 4L flow cytometer. JC-1 green and red fluorescence were detected in the FITC and PE channels, respectively.

Flow cytometry data were analyzed using FlowJo version 10. Mitochondrial membrane potential was quantified as the ratio of the geometric mean fluorescence intensity of JC- 1 red fluorescence to JC-1 green fluorescence (PE/FITC ratio) in the bulk cell population. A lower PE/FITC ratio was interpreted as reduced mitochondrial membrane potential and increased mitochondrial depolarization.

## Western Blotting

Cells were washed with 1× phosphate-buffered saline (PBS) and lysed in radioimmunoprecipitation assay (RIPA) buffer supplemented with protease inhibitor and phosphatase inhibitor. Lysates were incubated on ice for 20 min and subsequently sonicated at 100 amp using alternating 5-s pulses and 5-s rest intervals for 5 minutes total duration. Lysates were then centrifuged at 13,000 × *g* for 5 min at 4 °C, and the resulting supernatants were collected. Total protein concentration was determined using a Bradford protein assay.

Equal amounts of protein (15-25 μg per sample) were mixed. For immunoblotting with the Total OXPHOS antibody cocktail, samples were incubated at 37 °C for 30 min prior to electrophoresis. For all other protein targets, samples were denatured at 95 °C for 5 min. Proteins were separated by sulfate polyacrylamide gel electrophoresis (SDS– PAGE) at 100 V for approximately 1.5 h using Biorad 4-12% SDS gel and transferred onto polyvinylidene difluoride (PVDF) membranes using a Bio-Rad wet-transfer system under 100 V for 1 hour.

Following transfer, membranes were blocked for 1 h at room temperature in 5% non-fat dry milk prepared in Tris-buffered saline containing 0.05% Tween-20 (TBST).

Membranes were incubated overnight at 4 °C with gentle rocking with primary antibodies diluted in 5% milk/TBST. Primary antibodies included WT1, PDK4, TOMM20, and the Total OXPHOS antibody cocktail, which detects NDUFB8 (Complex I), SDHB (Complex II), UQCRC2 (Complex III), MT-CO2/COXII (Complex IV), and ATP5A (Complex V). Vinculin was used as a loading control where applicable.

Following primary antibody incubation, membranes were washed three times with TBST and incubated for 1 h at room temperature with the appropriate horseradish peroxidase (HRP)-conjugated anti-rabbit or anti-mouse secondary antibody. Membranes were subsequently washed three times with TBST. Chemiluminescent signal was developed using SuperSignal™ West Femto Maximum Sensitivity Substrate by combining the two substrate components at a 1:1 ratio and incubating membranes for approximately 15 s to 1 min before imaging. Chemiluminescence was detected using a Bio-Rad ChemiDoc imaging system.

Densitometric analysis of immunoblot bands was performed using ImageJ/Fiji. Band intensities for WT1 and PDK4 were normalized to vinculin as a loading control, whereas individual OXPHOS complex protein signals were normalized to TOMM20 to account for mitochondrial content. Normalized protein expression was expressed as fold change relative to the corresponding nMSC, empty vector, or scramble control, as applicable.

## Cancer stimulation and tumor microenvironmental stress and mitochondrial transfer assays

To assess the effects of cancer stimulation and tumor microenvironmental stress on mitochondrial oxidation, nMSC expressing mitochondrial matrix-targeted HyPer7 (mitoHyPer7) were cultured for five days under adherent or ultra-low attachment (ULA) conditions, either alone or in coculture with OVCAR3 ovarian cancer cells. For adherent conditions, cells were seeded in six-well tissue culture-treated plates at 30,000 nMSC per well for monoculture or 15,000 nMSC and 15,000 OVCAR3 cells per well for coculture. Cells were maintained in a 1:1 mixture of complete MSC medium and OVCAR3 culture medium, as described above. For ULA conditions, cells were seeded in six-well ULA plates at 50,000 nMSC per well for monoculture or 25,000 nMSC and 25,000 OVCAR3 cells per well for coculture. ULA cultures were maintained in serum- free MEBM containing the same supplements as complete MSC medium, excluding fetal bovine serum. Corresponding nMSC-only controls were maintained in the same culture medium as their respective coculture conditions.

To distinguish MSC from ovarian cancer cells during flow cytometric analysis, OVCAR3 cells were labeled with CellTrace Far Red (CTFR; Invitrogen) prior to coculture. CTFR stock solution was prepared according to the manufacturer’s instructions and diluted 1:1000 in phosphate-buffered saline (PBS). Tumor cells were incubated with CTFR for 30 min, followed by quenching in serum-containing culture medium for 5 min and washing twice before coculture. Following five days of culture, cells were harvested by trypsinization, pelleted, and resuspended in ice-cold flow cytometry buffer. Samples were maintained on ice until analysis. Following singlet gating, mitochondrial oxidation was measured specifically within the CTFR-negative, mitoHyPer7-positive MSC population using the 488/405 fluorescence ratio.

To assess the oxidative state of mitochondria transferred from CA-MSC to ovarian cancer cells, mitoHyPer7-expressing CA-MSC were cocultured with CTFR-labeled OVCAR3, CAOV3, or OVSAHO cells in six-well ULA plates. CA-MSC and tumor cells were seeded at a 1:1 ratio (25,000 cells of each type per well) in serum-free supplemented MEBM and cocultured for 24 h. Following coculture, cells were harvested and analyzed by flow cytometry. After singlet gating, CTFR-negative, mitoHyPer7- positive cells were identified as donor CA-MSC, whereas CTFR-positive, mitoHyPer7- positive cells were identified as recipient tumor cells containing CA-MSC-derived mitochondrial signal.

For population-level analysis, mitochondrial oxidation was quantified by dividing the geometric mean fluorescence intensity following 488-nm excitation by that following 405-nm excitation within each gated population. For single-cell analysis, fluorescence intensities of individual events were exported from FlowJo into Microsoft Excel, where the 488/405 fluorescence ratio was calculated separately for each cell. These ratios were used to compare the oxidation state of CA-MSC-derived mitochondrial signal detected within recipient tumor cells with that retained in donor CA-MSC.

## Live cell timelapse imaging

Live-cell time-lapse imaging was performed to visualize mitochondrial transfer from CA- MSC to ovarian cancer cells and assess the oxidative state of mitochondria during transfer. OVCAR3 cells labeled with CellTrace Far Red (CTFR) were seeded at 2,000 cells per MatTek 60-mm glass-bottom dish in complete OVCAR3 culture medium. The following day, 30 min before imaging, the medium was replaced with a 1:1 mixture of MSC and OVCAR3 culture media, each supplemented with 2% fetal bovine serum, and 2,000 mitoHyPer7-expressing CA-MSC were added.

Live-cell imaging was performed using a Leica confocal microscope equipped with a 20× objective and maintained at 37 °C with 5% CO₂. Time-lapse images were acquired at 15-min intervals over 24 h, with 12 optical Z-sections collected at approximately 0.69- µm intervals. Three fluorescence channels were acquired to visualize CTFR-labeled tumor cells and mitoHyPer7 fluorescence following excitation at 488 nm and 405 nm.

Images were processed using Fiji/ImageJ, and mitochondrial oxidation was visualized using the mitoHyPer7 488/405 fluorescence ratio. Individual Z-sections were examined to assess heterogeneity in mitochondrial oxidation within donor CA-MSC. Mitochondrial transfer events were qualitatively tracked over time, with optical sections manually selected across successive time points to follow the mitochondrial signal through the Z- stack. Representative time-lapse images and the corresponding supplementary video were prepared to visualize mitochondrial oxidation before, during, and following transfer into recipient tumor cells.

## Mitochondrial Single-Cell Assay for Transposase Accessible Chromatin with sequencing (mtscATAC-seq)

### Nuclei preparation

Nuclei and mitochondria from N=3 nMSC and N=3 CA-MSC donors were isolated as previously described by Lareau et al. (2023)^51^ -- with modification to improve nuclear integrity and yield. Per donor, 200,000 cryopreserved cells were rapidly thawed at 37°C and diluted in 1mL 50% FBS Flow Buffer (v/v) (HBSS, 25mM HEPES, heat-inactivated FBS, 0.22µm filtered). Cellular suspensions were further diluted stepwise to 50mL using 2% FBS Flow Buffer (v/v) (HBSS, 25mM HEPES, heat-inactivated FBS, 0.22µm filtered). Diluted cell suspensions were pelleted (300xg, 10 min, 4°C), and supernatant was removed. The cells were then washed in 2% FBS Flow Buffer, pelleted (300xg, 5 min, 4°C), and the supernatant was removed. Next, the cells were resuspended in 2% FBS Flow Buffer and counted using fluorescence-based Acridine Orange/Propidium Iodide (AO/PI; 1:10 final dilution) on a Luna-FX7 Automated Cell Counter. This input allowed for roughly 50% cell loss between thawing and final nuclei count.

Cells were lightly fixed in 1% (v/v) formaldehyde (10 min, room temperature) and quenched with glycine solution (0.125 M final concentration). Next, the cells were washed twice -- first with pH 7.4 PBS, then with 2% FBS Flow Buffer. Between washes, cells were pelleted (400xg, 5 min, 4°C, swinging bucket rotor), and the supernatant was removed.

The cells were permeabilized for 3 min, on ice, in chilled lysis buffer (1x Resuspension Buffer [10 mM Tris-HCl pH 7.4, 10 mM NaCl, 3 mM MgCl2] with 0.1% NP-40 and 1% bovine serum albumin (BSA)). Cells were then washed with wash buffer (1x Resuspension Buffer with 1% BSA) and pelleted (500xg, 5 min, 4°C, swinging bucket rotor). Using a prewetted pipette tip, the supernatant was removed gently without disrupting the pellet.

Fixed, lysed cells were resuspended slowly in chilled 1x diluted nuclei buffer (10x Genomics) at ∼4,000 cells/µL. Fixed, lysed cells were counted with AO/PI, as described above. Because permeabilized nuclei take up PI, the PI-positive count was used as the final nuclei count.

### Library Construction

Libraries were generated using the *Chromium Next GEM Single Cell ATAC Reagent Kits v2* (10x Genomics; User Guide CG000496 – Rev B), with modifications as previously described^51,52^. Tagmentation was extended to 60 minutes (Step 1.2), and one additional PCR cycle was performed during sample index polymerase chain reaction (PCR) (Step 4.1). Final libraries were eluted in EB buffer and stored in DNA LoBind tubes at -20°C. Libraries were quantified with the Qubit dsDNA HS Assay Kit, and fragment size distribution was assessed using an Agilent Bioanalyzer 2100.

### Sequencing & Quality Control

Single-cell ATAC-seq libraries underwent 2x150bp paired-end sequencing on the Element AVITI™ (Element Biosciences) at the Van Andel Institute Genomics Core. Base calling was performed on-instrument, and AVITI OS v2.6.2 output was demultiplexed and converted to FastQ format with **Bases2fastq** (v1.7.0, Element Biosciences). Read quality control was performed on the raw FASTQ files using **FastQC** v0.12.1 and **MultiQC** v1.27.1.

### Custom NUMT-masked Reference Preparation

To minimize off-target alignment of mitochondrial reads to nuclear mitochondrial DNA segments (NUMTs), a custom reference was built per the instructions of Lareau et al., 2023^51^. The human nuclear genome (GRCh38; 10x Genomics Single Cell ATAC Reference - 2020-A-2.0.0 (May 3. 2021) was hard-masked using a human mitochondrial blacklist bed file (hg38.full.blacklist.bed; downloaded from https://github.com/caleblareau/mitoblacklist/tree/master/combinedBlacklist) using **bedtools** (v2.31.1). Following masking of the nuclear genome, the final custom reference was generated using cellranger-atac mkref, using **Cell Ranger ATAC** (v2.1.0 compatible with Element AVITI base qualities; 10x Genomics).

### Read Alignment

Single-cell sequencing data were aligned to the custom reference using the cellranger- atac count function (v2.1.0, Element AVITI-compatible build; 10x Genomics), using default parameters^51^. Adapter sequences generated during Tn5-based tagmentation were trimmed internally by the count function prior to alignment. Across donors, both CA-MSC and NMSC had an average of 93.7% read pairs confidently mapped. Further, CA-MSC had an average of 3,894 cells at an average depth of 76,910 raw read pairs/cell, and NMSC had an average of 5,863 cells at an average depth of 46,972 raw read pairs/cell.

### Mitochondrial variant calling with Mitochondrial Genome Analysis Toolkit

Per-cell mitochondrial allele counts and variant detection occurred using Mitochondrial Genome Analysis Toolkit (**mgatk** v0.7.0), following Lareau et al., 2023^51,52^. mgatk was run in tenx mode using parameters -bt CB -g hg38. For every cell, mgatk counted each of the four bases on each strand of all 16,569 mtDNA positions and computed coverage, per-cell variant allele frequencies (VAF), and per-variant summary statistics. mgatk coverage output files were imported into **R** (v4.4.2) and annotated with sample identifiers. Each cell’s mean mtDNA coverage was taken from mgatk’s depth table (*.depthTable.txt). Following default mgatk parameters, only cells with mean mitochondrial coverage of at least 10x were used for downstream variant calling.

### Identification of high-quality variants

Mitochondrial variants were filtered per-sample using mgatk variant statistics output (*.variant_stats.tsv.gz). For each variant, VAF (heteroplasmic fraction) is defined by the number of reads supporting the variant allele divided by the sum of reads covering that position in a cell. A variant was considered high-quality if it met three criteria: (1) it was confidently detected in at least 3 cells -- meaning at least 2 forward-strand and 2 reverse-strand reads support it in at least 3 cells, (2) it had strand correlation greater than 65%, where strand correlation is Pearson correlation between forward- and reverse-strand variant read accounts across cells, and (3) its mean coverage at the variant position was at least 10x. Across 6 samples, 251 unique variants were confidently detected in at least 3 cells. Of these, 209 met all high-quality criteria (37-55 per sample).

All high-quality variants, including homoplasmic and heteroplasmic, were kept for downstream analysis. A variant was considered heteroplasmic if the variance-to-mean VAF ratio (VMR) across cells was greater than 0.01; otherwise, it was considered homoplasmic (≤ 0.1 VMR). A variant was considered present in a cell if its VAF was greater than 0.01. A variant reached homoplasmy with if at least one cell had VAF ≥ 0.95 for said variant.

### Mitochondrial variant calling in bulk ATAC-seq

Mitochondrial DNA variants were called from eight publicly-available bulk ATAC-seq libraries (n = 4 nMSC, n = 4 CA-MSC; accession number <u>SRP289005</u>)^21^. Variant calling followed the adapted GATK workflow described by Laricchia & Lake et al. (2022)^53^.

Reads were aligned with bwa mem (v0.7.19) to the NUMT-masked GRCh38 reference. Duplicates were marked with **samblaster** (v 0.1.26) and unmapped/low-quality reads were removed using **samtools** (v1.21, MAPQ < 20). Mitochondrial variants were called with **GATK** (v4.6.2.0) Mutect2 --mitochondria-mode. All mitochondrial-assigned fragments were then extracted and realigned to mitochondrial reference shifted by 8,000bp; this was to ensure detection of variants across the control region, which spans the necessary artificial breakpoint needed to linearize the mitochondrial genome during alignment. Shifted-alignment calls were then lifted back to standard mitochondrial coordinates using **Picard** (4.3.0) LiftoverVcf and merged with unshifted calls for each sample using **Picard** MergeVcfs. Calls were filtered with GATK FilterMutectCalls, masked against a blacklist of known artifact-prone sites, and multiallelic sites were split. Per-library VCFs were merged by condition via **bcftools** (v1.21), annotated against gnomAD v3.1 chrM allele frequencies, and variants exceeding 0.4% population frequency were removed. This process yielded a list of putatively somatic (population- rare) variants, since no matched normal was available.

### Variant Pathogenicity Annotation

Single nucleotide variants from single-cell and bulk ATAC-seq datasets were annotated using the MitImpact 4 annotation feature (mitimpact.mcb2lab.org/annotate). Protein- coding variants were classified using APOGEE2^54^, and tRNA/rRNA variants were classified using nAPOGEE^55^. The classification scale was based on a seven-tier scale: Pathogenic, likely pathogenic, VUS+, VUS, VUS-, likely benign, and benign. Control region variants were not annotated. For condition-level summaries, each unique variant was counted once per condition, using the most severe class and highest heteroplasmy detected.

## Isolation of MSC Mitochondria

Mitochondria were isolated from MSCs and CA-MSCs via differential centrifugation using the mitochondria Isolation Kit for Cultured Cells per manufacturer instructions (Abcam). Following kit isolation, mitochondria were resuspended in 60µL 1X MAS buffer (220 mM of d-Mannitol, 70 mM of sucrose, 10 mM of KH2PO4, 5 mM of MgCl2, 2 mM of HEPES, 1 mM of EGTA, and 0.2% (w/v) of fatty acid-free BSA in ultrapure H2O and adjust the pH to 7.2 with KOH at 37 °C) and kept on ice until use. Mitochondrial yield was determined with the Pierce BCA Protein Assay Kit (Fisher Scientific). Mitochondria integrity was assessed by staining seeded mitochondria with 50nM MitoTracker Deep Red (Fisher Scientific) for 30 minutes, gently washing with fresh 1X MAS buffer, and imaging using fluorescent microscopy.

## Isolated Mitochondrial Complex Seahorse Assay

To assess the bioenergetic function of the individual mitochondria complexes, 10µg of isolated MSC/CA-MSC mitochondria in a total volume of 50uL was loaded into 3 wells of a Seahorse XFp plate and centrifuged at 2,200xg for 20 minutes at 4°C. While spinning, the cartridge ports were loaded with the appropriate compounds. For Complex I or II assays, port A was loaded with 20µL of ADP (40µM), port B with 22µL oligomycin (25µg/mL), port C with 24µL FCCP and port D with 26µL rotenone (20µM). For complex III/IV assays, port A was loaded with 20µL of ADP (40µM) and duroquinol (50mM), port B with 22µL oligomycin (25µg/mL) and port C with 24µL potassium azide (200mM).

Once centrifuged down, 130µL of prewarmed 1X MAS buffer containing the appropriate substrate (Complex I: 10mM glutamate and 5mM malate; Complex II: 10mM succinate and 2µM rotenone; Complex III/IV: 1X MAS only) was gently added to each well. OCR was measured on the Seahorse XFp platform and data analyzed as previously reported^56^.

## Bulk RNA-seq

Patient-derived nMSCs and CA-MSCs were cultured as previously described until reaching 70% confluency, pelleted and total RNA isolation carried out using a Nucleospin RNA Isolation Kit (Takara Bio USA, USA). RNA concentration and quality was assessed via a NanoDrop spectrophotometer. Samples were then packaged and shipped to Novogene Corporation for further processing and sequencing. mRNA was purified from total RNA using poly-T oligo-attached magnetic beads. After fragmentation, first-strand cDNA synthesis was carried out using random hexamer primers followed by second strand synthesis using dUTP. Qubit (Thermo Fisher Scientific, Waltham, MA) and real-time PCR was used for quantification and bioanalyzer for size distribution detection. Quantified libraries were pooled and sequenced on an Illumina platform (Illumina Inc., San Diego, CA) and 150-bp paired-end reads were generated.

Raw reads in fastq format were processed through a Novogene in-house perl script for QC. Briefly, reads containing adapters, ploy-N, and low-quality reads were removed and Q20, Q30, and GC content was calculated. The index of the reference genome hg38 was built and paired-end clean reads were aligned to the reference genome using hisat2 (v2.0.5). Quantification of the gene expression level was done via featureCounts (v1.5.0-p3). The FPKM of each gene was calculated based on the length of the gene and read counts mapped to the gene. Differential gene expression analysis was carried out using R (v4.1.3) package DESeq2 (v1.20.0). Functional enrichment analysis was done using the R package clusterProfiler and Gene Set Enrichment Analysis (GSEA)(Broad Institute, USA).

## Patient tumor single-cell RNA-seq

### Patient tumor single-cell RNA-seq processing and sorting

Fresh ovarian tumor tissue was obtained from the operating room (OR) from a treatment-naive patient with ovarian cancer and processed into a single-cell suspension within approximately 1 h of collection. Briefly, tumor tissue was mechanically dissected into small pieces and enzymatically digested with a mixture of collagenase A and hyaluronidase for 1 h at 37 °C. Following digestion, the cell suspension was passed through a 70-μm cell strainer to remove undissociated tissue and debris.

The resulting single-cell suspension was used to generate three cell populations for downstream single-cell RNA sequencing (scRNA-seq): freshly dissociated bulk tumor cells, cultured tumor cells, and cultured carcinoma-associated mesenchymal stem cells (CA-MSCs). For the cultured tumor cell population, cells were maintained in DMEM/F12 medium supplemented with 10% fetal bovine serum (FBS) and 1% penicillin– streptomycin. Tumor cell identity was validated by EpCAM staining. For the cultured CA- MSC population, cells were maintained in complete MSC medium as described above. CA-MSC identity was validated by flow cytometric analysis of the cell-surface markers CD105, CD90, and CD73.

On the day of sorting, all three cell populations were stained with DAPI for live/dead cell discrimination and with sample-hashing antibodies to allow the different populations to be sorted together while retaining their sample identities.

Hashed patient bulk ovarian tumor cells, cultured CA-MSCs, and cultured tumor cells were sorted directly onto the 10X chip using the ThermoFisher Scientific Bigfoot Spectral Cell Sorter to maximize cell yield and quality for scRNA-seq.

### Single-cell suspension and GEM generation

Single-cell 5’ gene expression and cell surface protein libraries were generated with the Chromium GEM-X Single Cell 5’ Reagent Kits v3 with Feature Barcode technology for Cell Surface Protein. Cell suspensions were combined with the reverse transcription master mix (RT Reagent E, Poly-dT RT Primer B, Reducing Agent B, and RT Enzyme E) and PBS, targeting recovery of 20,000 cells per reaction. Each mixture was loaded onto a GEM-X Single Cell 5’ Chip with Single Cell 5’ v3 Gel Beads and Partitioning Oil B. GEMs were generated on a Chromium X using the GEM-X 5P program. Within each GEM, the cell was lysed and polyadenylated mRNA and the antibody-derived feature barcode oligonucleotides were barcoded with a 10x cell barcode and unique molecular identifier (UMI). GEMs were transferred to a tube strip and incubated for reverse transcription.<u>Post-GEM-RT cleanup and cDNA amplification</u>

GEMs were broken with Recovery Agent and barcoded first-strand cDNA was purified with Dynabeads MyOne SILANE and eluted. cDNA and antibody-derived feature barcode products were amplified together by PCR with Amp Mix and Feature cDNA Primers. Amplified product was then separated by size with SPRIselect. A 0.6X selection left the larger cDNA from polyadenylated mRNA on the beads, which was washed and eluted for 5’ gene expression library construction. The supernatant containing the smaller antibody-derived products was retained and cleaned with a further 2.0X SPRIselect step for cell surface protein library construction. Twenty-five percent of the total cDNA yield was carried forward into gene expression library construction.

### 5’ gene expression library construction

Purified cDNA was fragmented, end-repaired, and A-tailed. The product underwent double-sided SPRIselect size selection (0.6X, then 0.8X), followed by adaptor ligation and a 0.8X SPRIselect cleanup. Libraries were indexed by sample index PCR with Amp Mix and a Dual Index TT Set A index. Libraries were cleaned by a second double-sided SPRIselect selection (0.6X, then 0.8X). Library size was assessed on an Agilent Bioanalyzer High Sensitivity chip and libraries were quantified.

### Cell surface protein library construction

Five microliters of the cleaned antibody-derived product were indexed by PCR with Amp Mix and a Dual Index TN Set A index. Products were purified with a 1.2X SPRIselect cleanup. Library size was assessed on an Agilent Bioanalyzer High Sensitivity chip and quantified.

### Sequencing

Gene expression and cell surface protein libraries were pooled and sequenced using paired-end dual-index reads at a recommended depth of at least 20,000 read pairs per cell for gene expression and 5,000 read pairs per cell for cell surface protein libraries.

## InferCNV

### Inference of copy number alterations from scRNA-seq data

Large-scale chromosomal copy number alterations were inferred from single-cell RNA- seq data with inferCNV (version 1.28.0). The input was the raw count matrix (RNA assay, counts layer) from the integrated, annotated Seurat object (version 5.5.0). Each cell was labeled with its final cell type annotation, and genes were ordered by genomic position using the GENCODE v27 annotation on the hg38 genome. T cells, B cells, and macrophages were designated as the reference populations. All other annotated populations were treated as observations. inferCNV was run with cutoff = 0.1, the value recommended for 10x Genomics data, so genes with a mean count below 0.1 across all cells were excluded. Reference-subtracted expression was smoothed along each chromosome and denoised (denoise = TRUE). Cells were clustered within their annotated groups (cluster_by_groups = TRUE), and the remaining parameters were left at their defaults.

### Chromosome-level summaries and clustering

Chromosome-level CNV signal was calculated per cell as the mean denoised relative expression across all genes on each chromosome and displayed as heatmaps (pheatmap, color scale 0.8 to 1.2). As an independent check on inferCNV’s built-in subclustering, cells were hierarchically clustered on the 2,000 most variable genes of the denoised matrix (Euclidean distance, Ward’s D2 linkage) and the tree was cut at *k* = 4. Agreement with the Leiden subclusters was assessed by cross-tabulation. Genome- wide heatmaps were redrawn with infercnv::plot_cnv (center = 1).

## Mitochondrial lineage tracing and phylogenetic reconstruction

Phylogenetic structure was inferred from mitochondrial DNA (mtDNA) sequence variation in single-cell RNA-sequencing data using Cellkin, a Python-based command- line pipeline. Reads aligning to the mitochondrial chromosome (chrM/MT) were extracted from a barcode-tagged BAM file against the GRCh38 reference genome FASTA. Extracted cells were restricted to a whitelist of cell barcodes of cells clustered in the tumor or stromal compartments and downsampled to a representative 2000 random cells for computational efficiency. A UMI-aware, per-cell pileup was then generated from the extracted reads. Candidate mtDNA variants were identified jointly across all cells from this pileup. Each cell was subsequently genotyped at the candidate sites, producing a variant allele frequency (VAF) and a discrete genotype call per cell per site. Prior to tree construction, cells were assessed for information content using per-cell quality metrics: total UMI depth, number of covered positions, number and fraction of variant sites with callable depth, mean variant depth, and mean variant VAF. Low- information cells were removed using thresholds on these metrics (--min-cell-total-umi- depth = 1000, --min-cell-called-variant-sites = 5000, --min-cell-called-variant-fraction = 0.2). Pairwise cell distances were computed from VAF profiles, by default as the depth- weighted mean absolute VAF difference over sites with sufficient depth in both cells.

Sites were first filtered to those well represented in the cohort, based on the fraction of cells with callable and the cohort-level VAF range. The pipeline also checks pairwise overlap, aborting if too large a fraction of cell pairs share no callable sites (--max-no- overlap-fraction = 0.2). Because of the size of the dataset (N = 2000 cells), distances were computed in Cellkin’s large-scale mode, which condenses pairwise distances (=distance-format = condensed, --large-scale-mode, --max-no-overlap-fraction = 0.2).

Pairwise mtDNA-based cell distances from Cellkin (condensed format) were analyzed in R together with the integrated Seurat object. Cells annotated as CAMSCs, MSCs, CAFs, or Tumor were retained to represent the tumor and stromal populations. To define mitochondrial clusters based on shared mt variants, the distance matrix was subjected to agglomerative hierarchical clustering (hclust, complete linkage), and the dendrogram was cut into k = 5 groups to represent transcriptional clusters. To relate mitochondrial lineage structure to transcriptional state, each cell pair was annotated with the integrated Seurat cluster of the first cell and the mitochondrial cluster of the second. For every combination of Seurat cluster and mitochondrial cluster, we calculated the mean pairwise distance, yielding a matrix of mean mtDNA distance between each transcriptional cluster and each mitochondrial cluster. Lower values indicate that cells in a transcriptional cluster are, on average, more closely related to the cells of that mitochondrial cluster. For visualization, the matrix was z-scored either by column, with values clipped to ±1.5. The scaled matrices were displayed as heatmaps (pheatmap), with Seurat clusters ordered by hierarchical clustering. Phylogenetic trees were taken from the hierarchal clustering done in this analysis and regenerated in Adobe Illustrator for clarity. *Software availability:* Cellkin is available at https://github.com/CilloLaboratory/Cellkin.

### Cluster 6 signature validation in external cohorts

A Cluster 6 transcriptional signature was generated from differential gene expression analysis comparing tumor Cluster 6 with the remaining tumor clusters (2, 7, 10, and 11) using the Seurat FindMarkers function with a Wilcoxon rank-sum test. Genes with an adjusted *P* value < 0.05 and average log2 fold change ≥ 0.5 were classified as Cluster 6-upregulated genes, whereas genes with an adjusted *P* value < 0.05 and average log2 fold change ≤ −0.5 were classified as Cluster 6-downregulated genes. The resulting signature consisted of 1,146 upregulated and 5,916 downregulated genes and was applied consistently across all independent validation cohorts.

Four independent bulk RNA-seq datasets containing patient-matched pre- and post- neoadjuvant chemotherapy (NACT) samples were analyzed: GSE143897 (n = 18 pairs), GSE227100 (n = 24 pairs), EGA HGSOC (n = 23 pairs), and Javellana HGSOC (n = 20 pairs). For GSE143897, Ensembl gene identifiers were converted to HGNC gene symbols using the org.Hs.eg.db R package, and counts corresponding to duplicate gene symbols were summed. For GSE227100, the deposited estimated-count matrix was used, with duplicate gene symbols similarly collapsed by summation. For both datasets, pre- and post-NACT samples were normalized together using the trimmed mean of M-values (TMM) method implemented in edgeR, followed by log2 counts-per- million (CPM) transformation with a prior count of 1.

For the EGA and Javellana cohorts, normalized reads per kilobase per million mapped reads (RPKM) expression matrices were obtained from the Townsend et al. HGSOC Gene Explorer^40^. Expression values were transformed as log2(RPKM + 1), and genes with missing expression values across all samples were excluded. These datasets were not subjected to additional TMM normalization because raw count matrices were unavailable.

Within each cohort, Cluster 6 signature genes represented in the corresponding expression matrix were retained, excluding genes with zero or non-finite variance. Gene expression values were standardized to Z-scores using the mean and standard deviation calculated across all matched pre- and post-NACT samples within each cohort. A Cluster 6-like signature score was then calculated for each sample by subtracting the mean Z-score of the downregulated signature genes from the mean Z- score of the upregulated signature genes. The two components were averaged separately to give equal weight to upregulated and downregulated genes, regardless of the number of genes in each group. No additional weighting based on fold change or statistical significance was applied.

Changes in Cluster 6-like signature scores were calculated for each patient as the difference between post-NACT and matched pre-NACT scores. Statistical significance was determined using a two-sided paired Wilcoxon signed-rank test. As one predefined signature comparison was performed per cohort, *P* values were not adjusted for multiple testing. Expression values were independently standardized within each cohort; therefore, absolute signature scores were not directly compared or pooled across datasets.

## Statistical Analysis

Statistical analyses were performed using GraphPad Prism (version 10.3.1) and R (version 4.4.2 for mitochondrial ATAC-seq analyses). Unless otherwise specified, comparisons between experimental groups were performed using two-sided, unpaired Student’s *t*-tests. Associations between WT1 protein expression and mitochondrial respiratory parameters were assessed using Pearson correlation analysis with simple linear regression. Comparisons of Cluster 6-like transcriptional signature scores between patient-matched pre- and post-neoadjuvant chemotherapy samples were performed using two-sided paired Wilcoxon signed-rank tests.

All statistical tests utilized for ATAC-seq data were performed in R (v4.4.2) using base R and the lme4/lmerTest packages, with Normal as the reference condition in every model. Bulk mtDNA variant burden was compared between conditions with a two-sided Wilcoxon rank-sum test (continuity-corrected). The distribution of unique bulk variants across mtDNA features was tested with Fisher’s exact test (simulated p-value, B = 10⁵, seed = 42). Per-cell mutational burden in single-cell data was modeled with a linear mixed-effects model on log₁₀(variants per cell + 1), with condition as a fixed effect and a random intercept for sample; the heteroplasmic-to-homoplasmic fraction was modeled with an analogous logistic mixed-effects model. Per-variant mean allele frequency distributions were compared between conditions with a two-sided Kolmogorov–Smirnov test. The proportion of pathogenic/likely-pathogenic variants reaching homoplasmy was compared between conditions with Fisher’s exact test. All tests were two-sided; p < 0.05 was considered significant.

More details on statistical analyses of mitochondrial DNA variants, differential gene expression, and pathway enrichment were performed as described in their respective Methods subsections where applicable.

Data are presented as mean ± standard error of the mean (s.e.m.), unless otherwise indicated. A *P* value < 0.05 was considered statistically significant. Exact *P* values are indicated in the corresponding figures where applicable.

## Experimental setup

For all in vitro experiments, a minimum of three independent experimental repeats were conducted unless otherwise indicated. Monoculture controls were used for coculture experiments where applicable. For sequencing and transcriptomic analyses, the number of independent patient samples and datasets analyzed is reported in the corresponding Methods/Results sections.

## Data availability

Publicly available datasets analyzed in this study are identified by their accession numbers in the corresponding Methods sections. Additional data generated during this study are available from the corresponding author upon reasonable request, subject to applicable ethical and data-sharing restrictions.

## Code availability

Custom code used for image processing and computational analyses is available from the corresponding author upon reasonable request.

## Acknowledgments

The authors thank the ProMark Gynecologic Oncology Biospecimen and Data Bank for facilitating access to patient-derived specimens. We acknowledge the University of Pittsburgh Center for Biologic Imaging (CBI) for providing microscopy resources and support for transmission electron microscopy and HyPer7 fluorescence imaging, and the UPMC Hillman Cancer Center Cytometry Facility (RRID:SCR_025361) for providing flow cytometry resources and services. We thank Dr. Wayne Stallaert and his laboratory for providing access to microscopy equipment for live-cell time-lapse imaging. We also thank the Delgoffe laboratory, particularly Rachel Cumberland, for facilitating access to and coordinating the use of the Seahorse Analyzer.

Work performed in the Pitt Biospecimen Core (RRID:SCR_025229) and services and instruments used in this project were supported, in part, by the University of Pittsburgh, the Office of the Senior Vice Chancellor for Health Sciences.

Research reported in this publication was supported by the National Cancer Institute of the National Institutes of Health under Award Number P30CA047904.

## References

1. Veneziani, A. C. et al. Heterogeneity and treatment landscape of ovarian carcinoma. Nat. Rev. Clin. Oncol. 20, 820–842 (2023).

2. Bowtell, D. D. et al. Rethinking ovarian cancer II: reducing mortality from high-grade serous ovarian cancer. Nat. Rev. Cancer 15, 668–679 (2015).

3. Gentric, G. et al. PML-regulated mitochondrial metabolism enhances chemosensitivity in human ovarian cancers. Cell Metab. 29, 156–173.e10 (2019).

4. Ghilardi, C. et al. PGC1α/β expression predicts therapeutic response to oxidative phosphorylation inhibition in ovarian cancer. Cancer Res. 82, 1423–1434 (2022).

5. Huang, Y. et al. Ct-OATP1B3 promotes high-grade serous ovarian cancer metastasis by regulation of fatty acid beta-oxidation and oxidative phosphorylation. Cell Death Dis. 13, 556 (2022).

6. Matassa, D. S. et al. Oxidative metabolism drives inflammation-induced platinum resistance in human ovarian cancer. Cell Death Differ. 23, 1542–1554 (2016).

7. Zampieri, L. X. et al. Mitochondria participate in chemoresistance to cisplatin in human ovarian cancer cells. Mol. Cancer Res. 18, 1379–1391 (2020).

8. Kleih, M. et al. Direct impact of cisplatin on mitochondria induces ROS production that dictates cell fate of ovarian cancer cells. Cell Death Dis. 10, 851 (2019).

9. Chandel, N. S. Mitochondria as signaling organelles. BMC Biol. 12, 34 (2014).

10. Chakrabarty, R. P. & Chandel, N. S. Mitochondria as signaling organelles control mammalian stem cell fate. Cell Stem Cell 28, 394–408 (2021).

11. Signorile, A. et al. Human ovarian cancer tissue exhibits increase of mitochondrial biogenesis and cristae remodeling. Cancers 11, 1350 (2019).

12. Song, M. et al. IRE1α-XBP1 controls T cell function in ovarian cancer by regulating mitochondrial activity. Nature 562, 423–428 (2018).

13. Xia, H., et al. Autophagic adaptation to oxidative stress alters peritoneal residential macrophage survival and ovarian cancer metastasis. JCI Insight 5, e141115 (2020).

14. Hwang, S. M. et al. Transgelin 2 guards T cell lipid metabolism and antitumour function. Nature 635, 1010–1018 (2024).

15. Atiya, H., Frisbie, L., Pressimone, C. & Coffman, L. Mesenchymal stem cells in the tumor microenvironment. Adv. Exp. Med. Biol. 1234, 31–42 (2020).

16. McLean, K. et al. Human ovarian carcinoma-associated mesenchymal stem cells regulate cancer stem cells and tumorigenesis via altered BMP production. J. Clin. Invest. 121, 3206–3219 (2011).

17. Coffman, L. G. et al. Ovarian carcinoma-associated mesenchymal stem cells arise from tissue-specific normal stroma. Stem Cells 37, 257–269 (2019).

18. McLean, K. et al. Leukemia inhibitory factor functions in parallel with interleukin-6 to promote ovarian cancer growth. Oncogene 38, 1576–1584 (2019).

19. Raghavan, S. et al. Carcinoma-associated mesenchymal stem cells promote chemoresistance in ovarian cancer stem cells via PDGF signaling. Cancers 12, 2063 (2020).

20. Cascio, S. et al. Cancer-associated MSC drive tumor immune exclusion and resistance to immunotherapy, which can be overcome by Hedgehog inhibition. Sci. Adv. 7, eabi5790 (2021).

21. Fan, H. et al. Epigenomic reprogramming toward mesenchymal-epithelial transition in ovarian-cancer-associated mesenchymal stem cells drives metastasis. Cell Rep. 33, 108473 (2020).

22. Frisbie, L. et al. Carcinoma-associated mesenchymal stem cells promote ovarian cancer heterogeneity and metastasis through mitochondrial transfer. Cell Rep. 43, 114551 (2024).

23. Goliwas, K. F. et al. Mitochondrial transfer from cancer-associated fibroblasts increases migration in aggressive breast cancer. J. Cell Sci. 136, jcs260419 (2023).

24. Sasaki, R., et al. Oxidative high mobility group box-1 accelerates mitochondrial transfer from mesenchymal stem cells to colorectal cancer cells providing cancer cell stemness. Int. J. Mol. Sci. 26, 1192 (2025).

25. Salaud, C. et al. Mitochondria transfer from tumor-activated stromal cells (TASC) to primary glioblastoma cells. Biochem. Biophys. Res. Commun. 533, 139–147 (2020).

26. Moschoi, R. et al. Protective mitochondrial transfer from bone marrow stromal cells to acute myeloid leukemic cells during chemotherapy. Blood 128, 253–264 (2016).

27. Echtay, K. S. et al. Superoxide activates mitochondrial uncoupling proteins. Nature 415, 96–99 (2002).

28. Talbot, D. A., Lambert, A. J. & Brand, M. D. Production of endogenous matrix superoxide from mitochondrial complex I leads to activation of uncoupling protein 3. FEBS Lett. 556, 111–115 (2004).

29. Ahmad, T. et al. Computational classification of mitochondrial shapes reflects stress and redox state. Cell Death Dis. 4, e461 (2013).

30. Yu, T., Sheu, S. S., Robotham, J. L. & Yoon, Y. Mitochondrial fission mediates high glucose-induced cell death through elevated production of reactive oxygen species. Cardiovasc. Res. 79, 341–351 (2008).

31. Pak, V. V. et al. Ultrasensitive genetically encoded indicator for hydrogen peroxide identifies roles for the oxidant in cell migration and mitochondrial function. Cell Metab. 31, 642–653.e6 (2020).

32. Castellana, S. et al. MitImpact 3: modeling the residue interaction network of the respiratory chain subunits. Nucleic Acids Res. 49, D1282–D1288 (2021).

33. Rath, S. et al. MitoCarta3.0: an updated mitochondrial proteome now with sub- organelle localization and pathway annotations. Nucleic Acids Res. 49, D1541–D1547 (2021).

34. Jiang, Y.-X. et al. Ascites-derived ALDH+CD44+ tumour cell subsets endow stemness, metastasis and metabolic switch via PDK4-mediated STAT3/AKT/NF-κB/IL-8 signalling in ovarian cancer. Br. J. Cancer 123, 275–287 (2020).

35. Dou, X. et al. PDK4-dependent hypercatabolism and lactate production of senescent cells promotes cancer malignancy. Nat. Metab. 5, 1887–1910 (2023).

36. Arend, R. C. et al. Metabolic alterations and WNT signaling impact immune response in HGSOC. Clin. Cancer Res. 28, 1433–1445 (2022).

37. Adzibolosu, N. et al. Immunological modifications following chemotherapy are associated with delayed recurrence of ovarian cancer. Front. Immunol. 14, 1204148 (2023).

38. Javellana, M. et al. Neoadjuvant chemotherapy induces genomic and transcriptomic changes in ovarian cancer. Cancer Res. 82, 169–176 (2022).

39. Pietilä, E. A. et al. Co-evolution of matrisome and adaptive adhesion dynamics drives ovarian cancer chemoresistance. Nat. Commun. 12, 3904 (2021).

40. Townsend, H. A. et al. Cross-assay RNA modeling reveals cancer biomarkers. Preprint at 10.64898/2026.04.30.722009 (2026).

41. Yakes, F. M. & Van Houten, B. Mitochondrial DNA damage is more extensive and persists longer than nuclear DNA damage in human cells following oxidative stress. Proc. Natl. Acad. Sci. USA 94, 514–519 (1997).

42. Vives-Bauza, C., Gonzalo, R., Manfredi, G., Garcia-Arumi, E. & Andreu, A. L. Enhanced ROS production and antioxidant defenses in cybrids harbouring mutations in mtDNA. Neurosci. Lett. 391, 136–141 (2006).

43. Tencerova, M. et al. Obesity-associated hypermetabolism and accelerated senescence of bone marrow stromal stem cells suggest a potential mechanism for bone fragility. Cell Rep. 27, 2050–2062.e6 (2019).

44. Bobba-Alves, N. et al. Cellular allostatic load is linked to increased energy expenditure and accelerated biological aging. Psychoneuroendocrinology 155, 106322 (2023).

45. Gao, Q. et al. Heterotypic CAF-tumor spheroids promote early peritoneal metastasis of ovarian cancer. J. Exp. Med. 216, 688–703 (2019).

46. Grieco, J. P., et al. Genetic and functional modifications associated with ovarian cancer cell aggregation and limited culture conditions. Int. J. Mol. Sci. 24, 14867 (2023).

47. Zhang, K. et al. Longitudinal single-cell RNA-seq analysis reveals stress-promoted chemoresistance in metastatic ovarian cancer. Sci. Adv. 8, eabm1831 (2022).

48. Croft, W. et al. The chromatin landscape of high-grade serous ovarian cancer metastasis identifies regulatory drivers in post-chemotherapy residual tumour cells. *Commun*. Biol. 7, 1211 (2024).

49. Garcia, G. L. et al. Aged and BRCA-mutated stromal cells drive epithelial cell transformation. Cancer Discov. 15, 1203–1224 (2025).

50. Chaudhry, A., Shi, R. & Luciani, D. S. A pipeline for multidimensional confocal analysis of mitochondrial morphology, function, and dynamics in pancreatic β-cells. Am. J. Physiol. Endocrinol. Metab. 318, E87–E101 (2020).

51. Lareau, C. A. et al. Mitochondrial single-cell ATAC-seq for high-throughput multi- omic detection of mitochondrial genotypes and chromatin accessibility. Nat. Protoc. 18, 1416–1440 (2023).

52. Lareau, C. A. et al. Massively parallel single-cell mitochondrial DNA genotyping and chromatin profiling. Nat. Biotechnol. 39, 451–461 (2021).

53. Laricchia, K. M. et al. Mitochondrial DNA variation across 56,434 individuals in gnomAD. Genome Res. 32, 569–582 (2022).

54. Bianco, S. D. et al. APOGEE 2: multi-layer machine-learning model for the interpretable prediction of mitochondrial missense variants. Nat. Commun. 14, 5058 (2023).

55. Bianco, S. D. et al. nAPOGEE: a machine-learning platform for clinically actionable pathogenicity assessment of all mitochondrial noncoding variants. Preprint at 10.64898/2025.12.20.695671 (2025).

56. Iuso, A., Repp, B., Biagosch, C., Terrile, C. & Prokisch, H. Assessing mitochondrial bioenergetics in isolated mitochondria from various mouse tissues using Seahorse XF96 analyzer. Methods Mol. Biol. 1567, 217–230 (2017).

