## Supplementary Information for "Stromal mitochondrial reprogramming and transfer reveal a chemotherapy-associated tumor state in ovarian cancer"

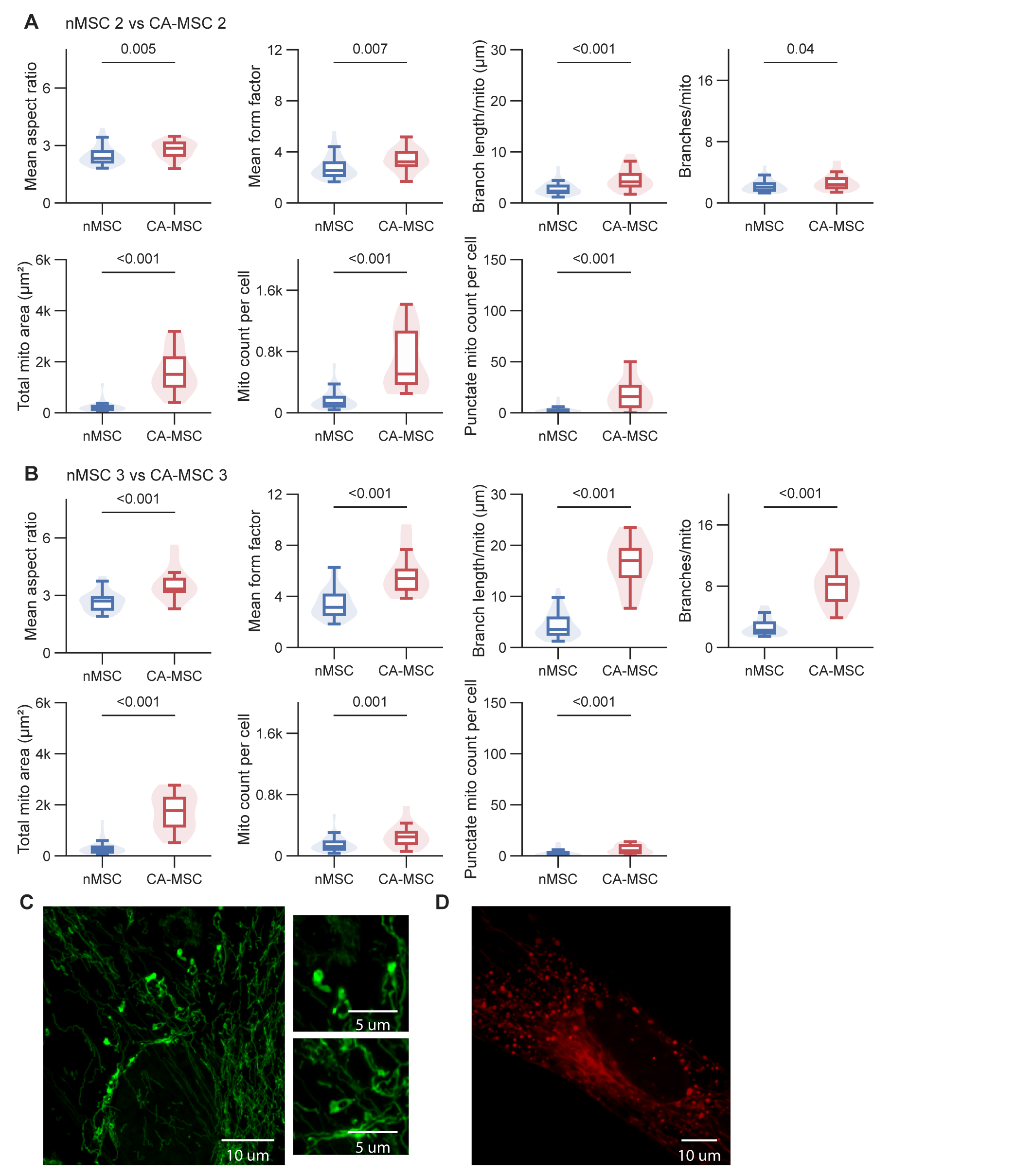

**Supplementary Figure 1 | Additional characterization of mitochondrial morphology in nMSC and CA-MSC.**

**A,B,** Quantification of mitochondrial morphology in independent patient-derived normal MSC (nMSC) and carcinoma-associated MSC (CA-MSC) lines, comparing nMSC2 with CA-MSC2 (A) and nMSC3 with CA-MSC3 (B). Measurements include mean aspect ratio, mean form factor, branch length per mitochondrion, branches per mitochondrion, total mitochondrial area per cell, mitochondrial count per cell, and punctate mitochondrial count per cell. **C,** Representative high-magnification (100×) confocal fluorescence images of COX8-labeled CA-MSC mitochondria showing distinct mitochondrial morphologies, including ring- and donut-shaped structures. Scale bars, 10 μm (left) and 5 μm (right). **D,** Representative confocal fluorescence image of CA-MSC mitochondria stained with MitoTracker Deep Red. Scale bar, 10 μm.

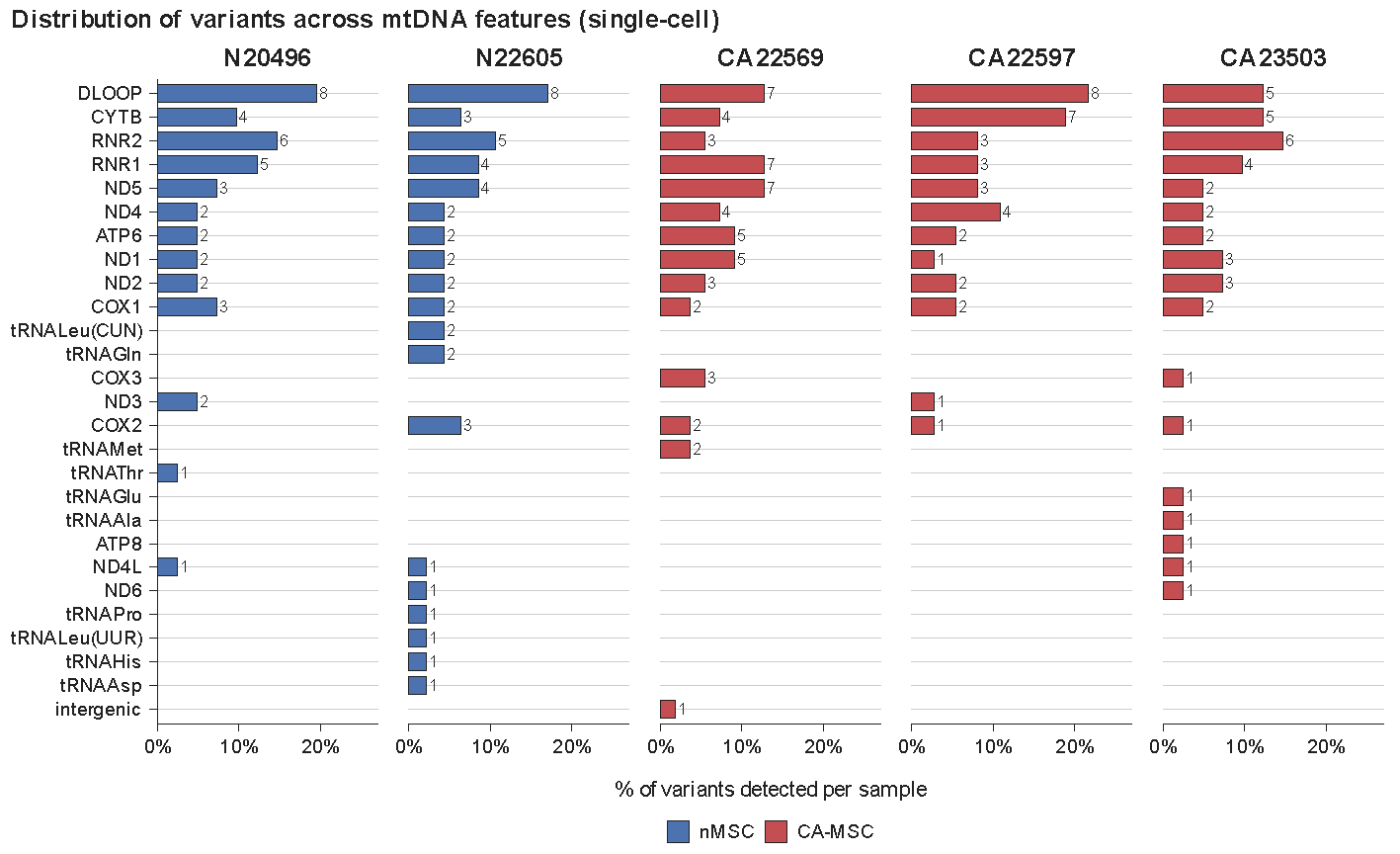

**Supplementary Figure 2 | Distribution of variants across the mitochondrial genome in single-cell, patient-derived MSC.**

Mitochondrial single-nucleotide variants detected in single-cell ATAC-seq data (N=3 CA-MSC, N=2 nMSC) using mgatk. Variant count per mitochondrial feature, per sample, is indicated to the right of each bar. Variant density across mtDNA features was assessed with a chi-square test of independence.

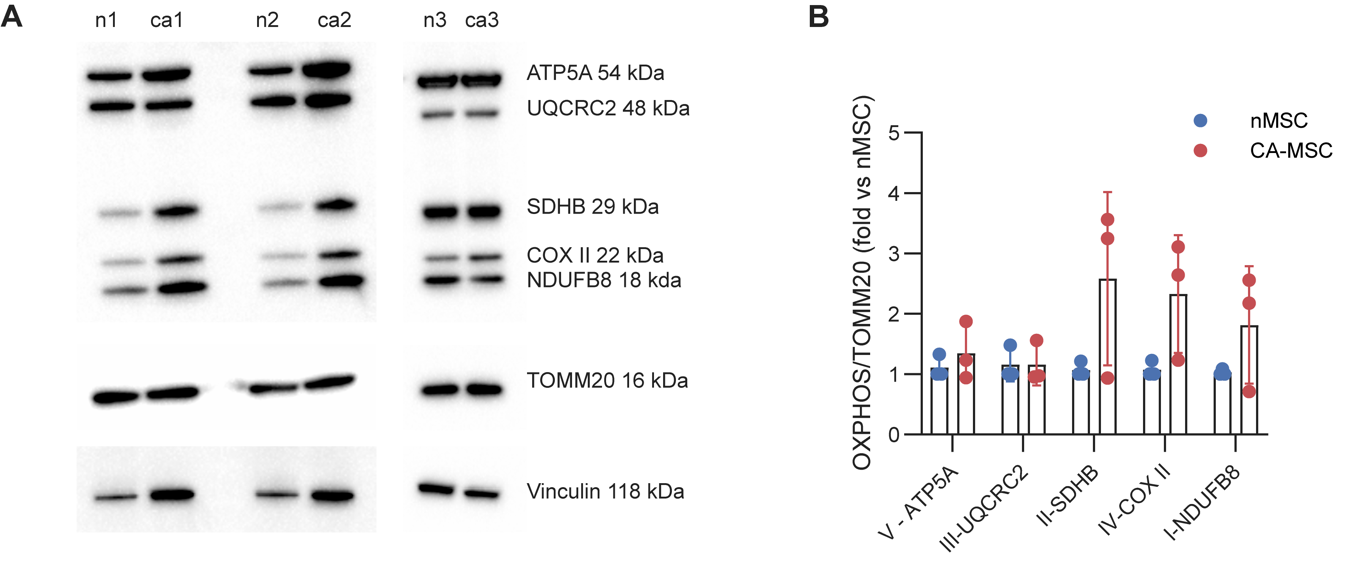

**Supplementary Figure 3 | Oxidative phosphorylation complex protein abundance in nMSC and CA-MSC.**

**A,** Immunoblot analysis of oxidative phosphorylation (OXPHOS) complex proteins in three independent patient-derived nMSC and three CA-MSC lines, using an antibody cocktail targeting representative subunits of mitochondrial respiratory complexes I–V. Two nMSC and three CA-MSC lines were also analyzed by mitochondrial single-cell ATAC sequencing (scATAC-seq). **B,** Densitometric quantification of individual OXPHOS complex proteins normalized to TOMM20 and expressed as fold change relative to the corresponding nMSC reference.

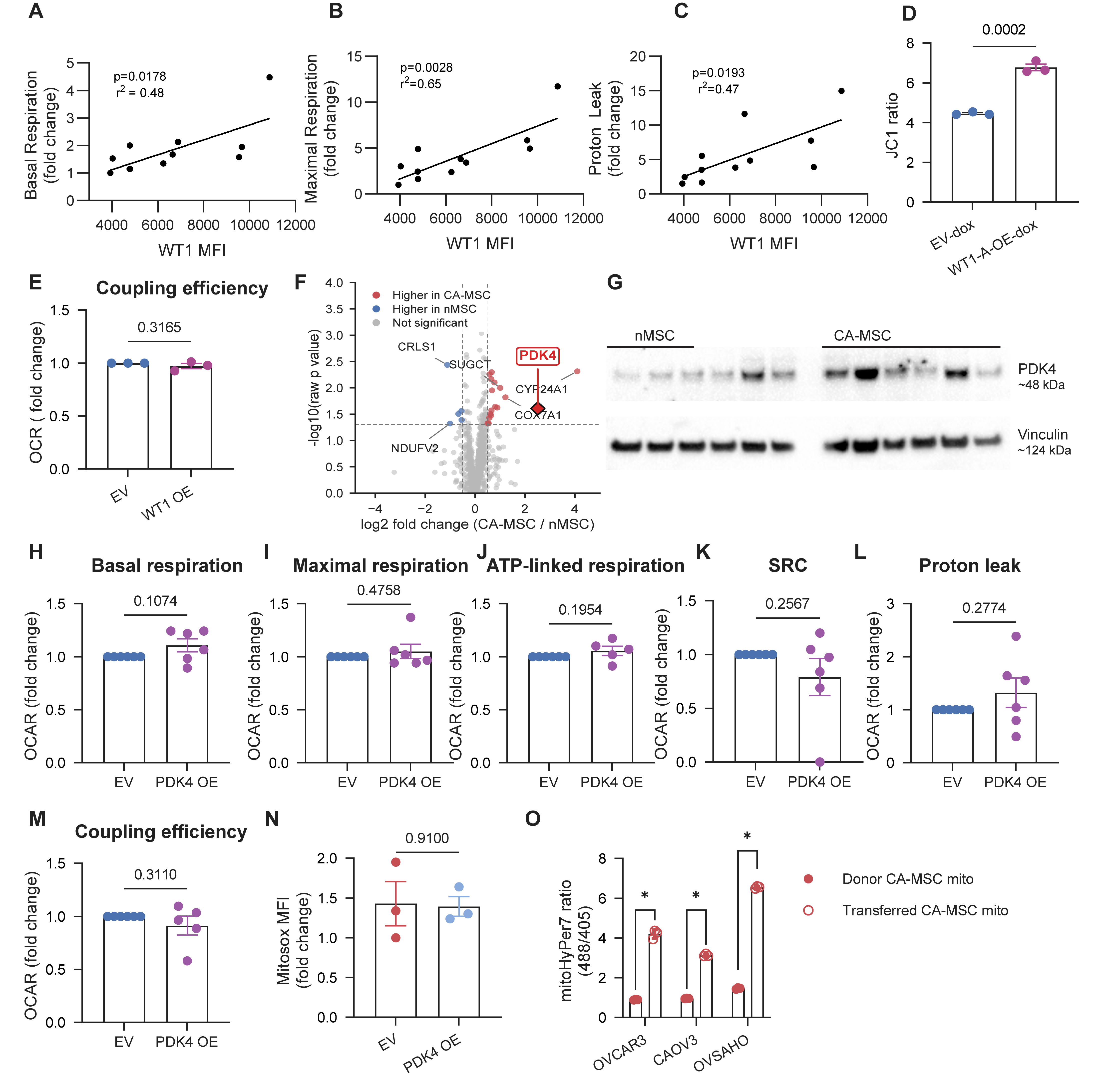

**Supplementary Figure 4 | Association of WT1 expression with mitochondrial respiration and additional characterization of WT1, PDK4, and mitochondrial transfer in MSCs.**

**A–C,** Pearson correlation analysis between WT1 protein expression, measured by flow cytometry, and basal respiration (A), maximal respiration (B), and proton leak (C) from Cell Mito Stress Tests across independent patient-derived MSC lines. Linear regression lines are shown, with coefficients of determination (R²) and corresponding *P* values indicated in the graphs. **D,** Flow cytometric quantification of mitochondrial membrane potential using JC-1 following doxycycline-inducible WT1 overexpression (OE) in normal MSC (nMSC) compared with empty vector (EV) controls. **E,** Quantification of mitochondrial coupling efficiency from Cell Mito Stress Tests following WT1 overexpression in nMSC. **F,** Volcano plot of differentially expressed MitoCarta genes identified by bulk RNA sequencing of patient-derived nMSC and carcinoma-associated MSC (CA-MSC), highlighting PDK4 among the upregulated genes in CA-MSC. **G,** Immunoblot analysis of PDK4 protein expression in three independent nMSC (first three lanes) and six independent CA-MSC lines (last six lanes), with vinculin as a loading control. **H–M,** Quantification of basal respiration (H), maximal respiration (I), ATP-linked respiration (J), spare respiratory capacity (K), proton leak (L), and coupling efficiency (M) from Cell Mito Stress Tests following PDK4 overexpression in nMSC. **N,** Flow cytometric quantification of mitochondrial superoxide using MitoSOX Red following PDK4 overexpression in nMSC. **O,** Quantification of mitochondrial HyPer7 oxidation ratios (488/405) in mitochondria retained within donor CA-MSC and CA-MSC mitochondria following transfer into OVCAR3, CAOV3, and OVSAHO ovarian cancer cells. Values represent geometric mean HyPer7 ratios calculated from three independent co-culture experiments for each tumor cell line.

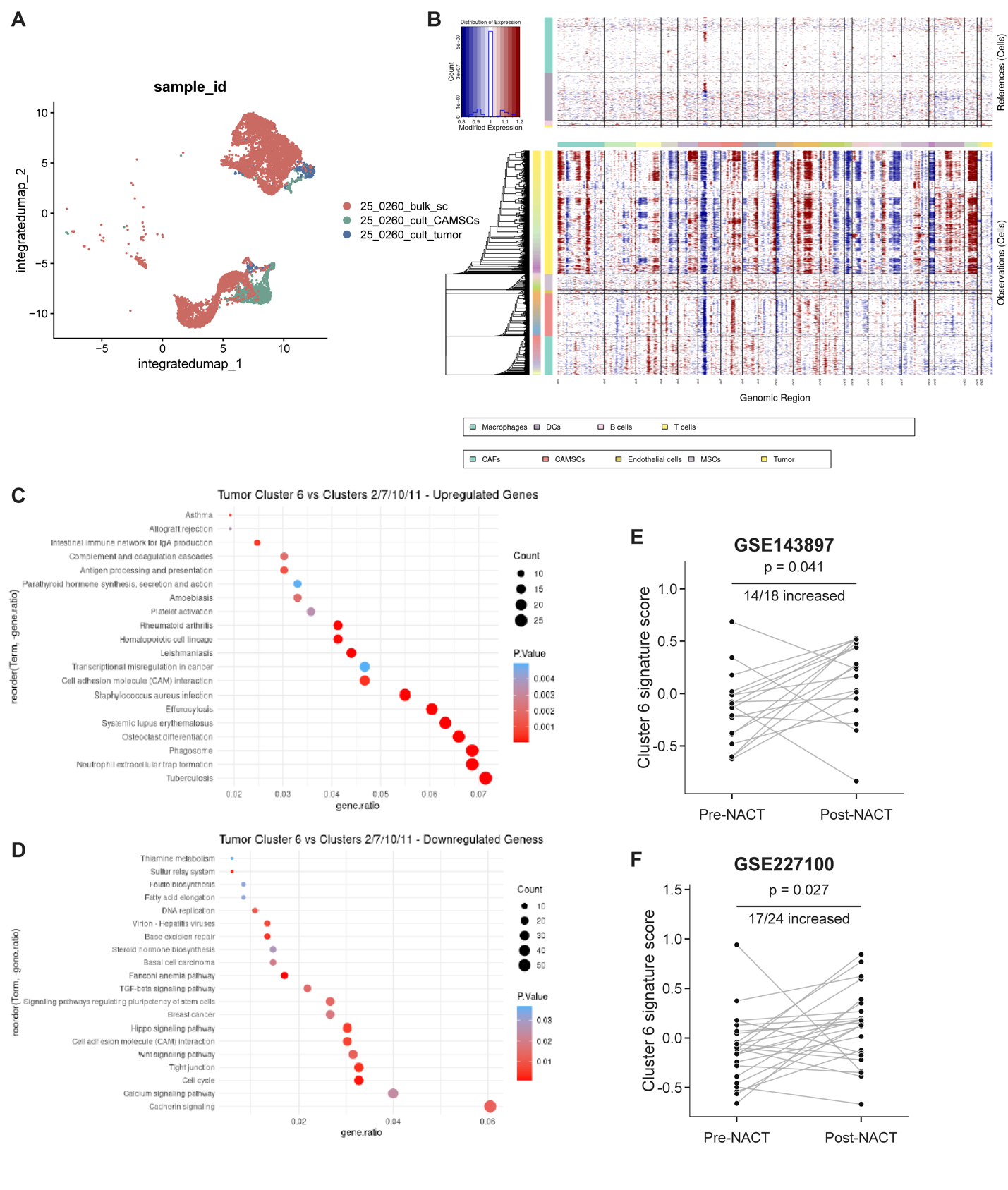
**Supplementary Figure 5 | Single-cell characterization of a distinct ovarian tumor cell population associated with CA-MSC mitochondrial lineage signatures.**

**A,** Uniform manifold approximation and projection (UMAP) of integrated single-cell RNA sequencing data from a patient-derived ovarian tumor specimen and matched cultured carcinoma-associated MSCs (CA-MSC) and tumor cells, colored by sample origin. **B,** Inferred copy-number variation (inferCNV) analysis of annotated cell populations using nonmalignant reference cells, showing chromosome-scale copy-number alterations in malignant tumor cells relative to stromal and immune populations. **C-D** Kyoto Encyclopedia of Genes and Genomes (KEGG) pathway enrichment analysis of genes upregulated (C) and downregulated (D) in tumor Cluster 6 compared with tumor Clusters 2, 7, 10, and 11. Dot size represents the number of genes associated with each pathway, and color indicates enrichment *P* values. **E,F,** Comparison of Cluster 6-like transcriptional signature scores between patient-matched pre-neoadjuvant chemotherapy (pre-NACT) tumors and post-NACT residual tumors from GSE143897 (E; n = 18 pairs) and GSE227100 (F; n = 24 pairs). Increased signature scores were observed following NACT in 14 of 18 patients in GSE143897 and 17 of 24 patients in GSE227100. Statistical significance was determined using two-sided paired Wilcoxon signed-rank tests (P = 0.0407 and P = 0.0268, respectively).

**Supplementary Table 1 | Clinical characteristics of MSC-derived cell lines**

| **Patient ID** | **Tissue site** | **MSC type** | **Age range** | **Histology** | **Neoadjuvant therapy** | **CRS** |
| --- | --- | --- | --- | --- | --- | --- |
| **nMSC** | | | | | | |
| 2045 | Fallopian tube | nMSC |  | Benign |  |  |
| 22605 | Fallopian tube | nMSC |  | Benign |  |  |
| 23505 | Fallopian tube | nMSC | 30–35 | Benign |  |  |
| 20496 | Fallopian tube | nMSC | 30–35 | Benign |  |  |
| 19327 | Fallopian tube | nMSC | 45–50 | Benign |  |  |
| **CA-MSC** | | | | | | |
| 20228 | Fallopian tube | CA-MSC | 75–80 | HGSOC | Yes | CRS3 |
| 23105 | Fallopian tube | CA-MSC |  | HGSOC | No |  |
| 23310 | Fallopian tube | CA-MSC | 60–65 | HGSOC | Yes | CRS2 |
| 22569 | Fallopian tube | CA-MSC | 80–90 | HGSOC | Yes |  |
| 23503 | Fallopian tube | CA-MSC | 55–60 | HGSOC | Yes | CRS3 |
| 19913 | Fallopian tube | CA-MSC | NA | HGSOC | Yes |  |
| 19609 | Fallopian tube | CA-MSC | 75–80 | HGSOC | Yes | CRS3 |
| 22597 | Fallopian tube | CA-MSC | 70–75 | HGSOC | Yes | CRS2 |
| 20254 | Fallopian tube | CA-MSC | 60–65 | HGSOC | Yes | CRS2 |

CRS = chemotherapy response score. CRS is not applicable to nMSC samples and is therefore left blank.

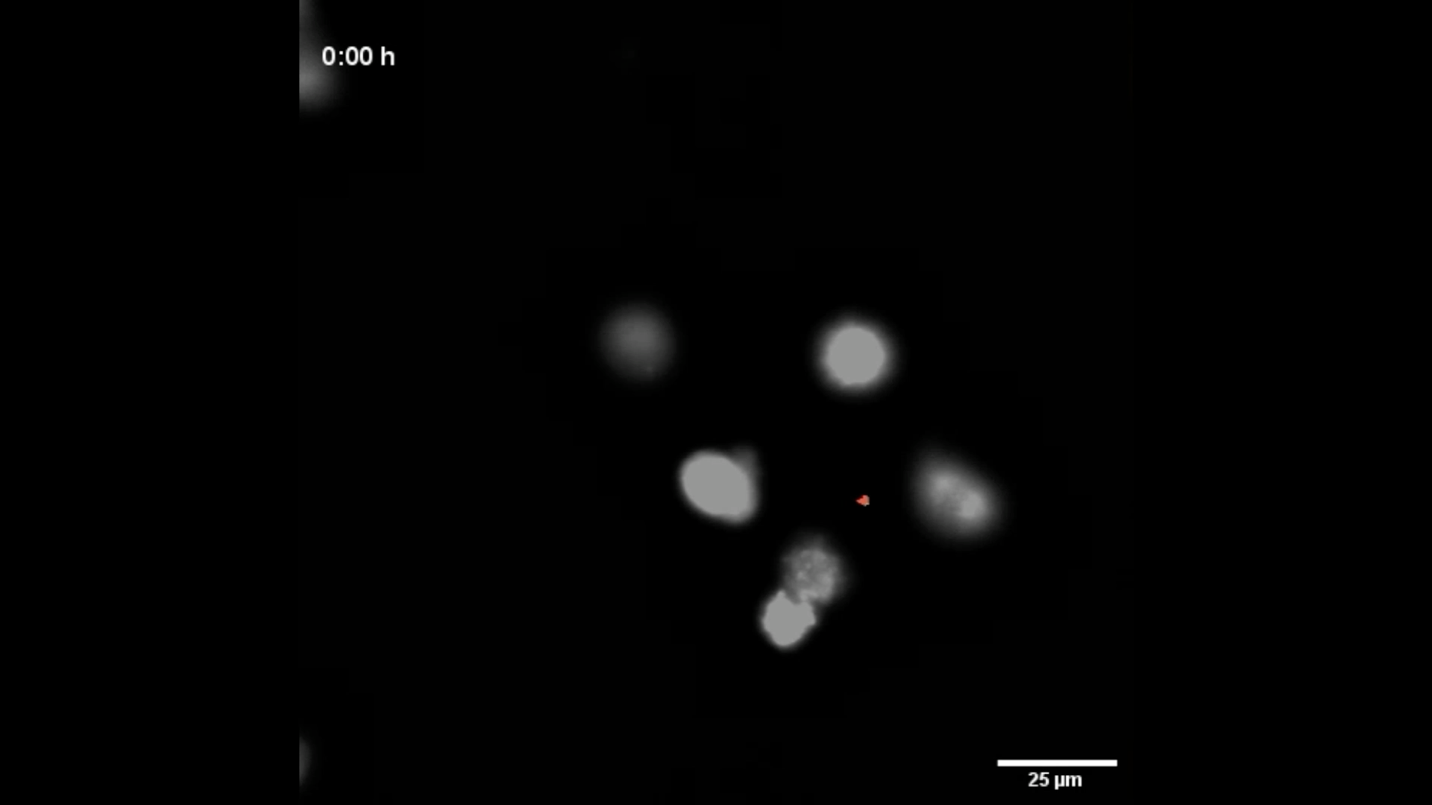
**Supplementary Video 1**

**Supplementary Video 1 | Time-lapse imaging of the transfer of oxidized mitochondria from CA-MSC to ovarian cancer cells.**

Representative time-lapse fluorescence microscopy showing mitochondrial transfer from carcinoma-associated MSC (CA-MSC) expressing mitochondrially targeted HyPer7 to CellTrace Far Red (CTFR)-labeled OVCAR3 ovarian cancer cells. The video displays pseudocolored ratiometric HyPer7 images generated from the 488/405 nm fluorescence signal, with red indicating the highest level of mitochondrial oxidation. The video captures CA-MSC approaching the tumor cell (0.5 h), establishing contact (3 h), transfer of CA-MSC mitochondrial signal into the recipient tumor cell (4.5 h), and persistence of the transferred mitochondrial signal within the recipient cell (8 h). The arrow indicates the transferred mitochondrial structure. Time is indicated in hours. Scale bar, 25 μm.
